# Functions of *Gtf2i* and *Gtf2ird1* in the developing brain: transcription, DNA-binding, and long term behavioral consequences

**DOI:** 10.1101/854851

**Authors:** Nathan D. Kopp, Kayla R. Nygaard, Katherine B. McCullough, Susan E. Maloney, Harrison W. Gabel, Joseph D. Dougherty

## Abstract

*Gtf2ird1* and *Gtf2i* may mediate aspects of the cognitive and behavioral phenotypes of Williams Syndrome (WS) – a microdeletion syndrome encompassing these transcription factors (TFs). Knockout mouse models of each TF show behavioral phenotypes. Here we identify their genomic binding sites in the developing brain, and test for additive effects of their mutation on transcription and behavior. Both TFs target constrained chromatin modifier and synaptic protein genes, including a significant number of ASD genes. They bind promoters, strongly overlap CTCF binding and TAD boundaries, and moderately overlap each other, suggesting epistatic effects. We used single and double mutants to test whether mutating both TFs will modify transcriptional and behavioral phenotypes of single *Gtf2ird1* mutants. Despite little difference in DNA-binding and transcriptome-wide expression, *Gtf2ird1* mutation caused balance, marble burying, and conditioned fear phenotypes. However, mutating *Gtf2i* in addition to *Gtf2ird1* did not further modify transcriptomic or most behavioral phenotypes, suggesting *Gtf2ird1* mutation alone is sufficient.

## Introduction

The Williams syndrome critical region (WSCR) contains 28 genes that are typically deleted in Williams syndrome (WS) (OMIM#194050). The genes in this region are of interest for their potential to contribute to the unique physical, cognitive, and behavioral phenotypes of WS, which include craniofacial dysmorphology, mild to severe intellectual disability, poor visuospatial cognition, balance and coordination problems, and a characteristic hypersocial personality (1–3). Single gene knockout mouse models exist for many of the genes in the region, with differing degrees of face validity for WS phenotypes (4–9). Two genes have been highlighted in the human and mouse literature as playing a large role in the social and cognitive tasks: *Gtf2i* and *Gtf2ird1*. Mouse models of each gene have shown social phenotypes as well as balance and anxiety phenotypes (4,8–12). Since evidence shows that each gene affects similar behaviors, we set out to test the hypothesis that simultaneous knockdown of both genes would cause more severe phenotypes. Investigating the genes together, rather than individually, could provide a more complete understanding of how the genes in the WSCR contribute to the phenotypes of WS.

*Gtf2i* and *Gtf2ird1* are part of the General Transcription Factor 2i family of genes. A third member, *Gtf2ird2*, is located in the WSCR that is variably deleted in WS patients with larger deletions (13). This gene family arose from gene duplication events, resulting in high sequence homology between the genes (14). The defining feature of this gene family is the presence of the helix-loop-helix I repeats, which are involved in DNA and protein binding (15). *Gtf2i*’s roles include regulating transcriptional activity in the nucleus. However, this multifunctional transcription factor also exists in the cytoplasm where it conveys messages from extracellular stimuli and regulates calcium entry into the cell (16, 17). So far, *Gtf2ird1* has only been described in the nucleus of cells and is thought to regulate transcription and associate with chromatin modifiers (18). The DNA binding of these two transcription factors (TFs) has only been studied in ES cells and embryonic craniofacial tissue. They recognize similar and disparate genomic loci, suggesting the proteins interact to regulate specific regions of the genome (19, 20). However, binding of these TFs has not been studied in the developing brain, which could provide more relevant insight on how the General Transcription Factor 2i family contributes to cognitive and behavioral phenotypes.

We performed ChIP-seq on GTF2I and GTF2IRD1 in the developing mouse brain to define where these TFs bind and then tested the downstream consequences of disrupting this binding. We used the CRISPR/Cas9 system to make a mouse model with a mutation in *Gtf2ird1* alone and a mouse model with mutations in both *Gtf2i* and *Gtf2ird1* to test how adding a *Gtf2i* mutation modifies the effects of *Gtf2ird1* mutation. We showed the mutation in *Gtf2ird1* resulted in the production of an N-truncated protein that disrupts the binding of GTF2IRD1 at the *Gtf2ird1* promoter and deregulates the transcription of *Gtf2ird1*, moderately decreasing protein levels so homozygous mutants approximate levels expected from hemizygosity of the WSCR. While there are mild consequences of the mutation on genome-wide transcription, the mutant mouse exhibited clear balance, marble burying, and conditioned fear differences. Comparing the single gene mutant to the double mutant did not reveal more severe transcriptional changes and behavioral phenotypes, however adding *Gtf2i* on top of a background of two *Gtf2ird1* mutations resulted in abnormal pre-pulse inhibition and cued fear conditioning. This suggests *Gtf2ird1* drives the majority of the phenotypes observed in current studies, and total protein level, N-terminal truncation, or both have functional consequences on DNA-binding and behavior.

## Results

### *Gtf2i* and *Gtf2ird1* bind at active promoters and conserved sites

The paralogous transcription factors, GTF2I and GTF2IRD1, have been implicated in the behavioral phenotypes seen in humans with WS as well as mouse models (4,8,11,12,21,22). However, the underlying mechanisms by which the General Transcription Factor 2i family acts are not well understood. One approach to begin to identify how these TFs can regulate phenotypes is by identifying where they bind in the genome. This has been done in ES cells and embryonic facial tissue and revealed that both of these transcription factors bind to genes involved in craniofacial development (19). However, these are not relevant tissues when considering phenotypes related to brain development and subsequent behavior. To address this, we performed ChIP-seq for GTF2IRD1 and GTF2I in the developing brain at embryonic day 13.5 (E13.5), a time point when both of these proteins are highly expressed (23).

We identified 1,410 peaks that were enriched in the GTF2IRD1 IP samples compared to the input (**Supplemental Table 1**). The GTF2IRD1-bound regions were strikingly enriched in the promoters of genes and along the gene body, more so than would be expected by randomly sampling the genome (Figure 1A) (χ^2^ = 1537.8, d.f. =7, p < 2.2×10^-16^). The bound peaks were found mostly in H3K4me3 bound regions (OR=779.5, p<2.2×10^-16^ Fisher’s exact test (FET)), suggesting they are in active sites in the genome. While GTF2IRD1-bound regions were also enriched in repressed regions of the genome as defined by H3K27me3 marks (OR=4.090, p< 2.2×10^-16^ FET), only 11% of GTF2IRD1 peaks were found in H3K27me3 regions as opposed to 94% in H3K4me3 regions (Figure 1B), suggesting GTF2IRD1 may have more of a role in activation than repression.

**Figure 1:**
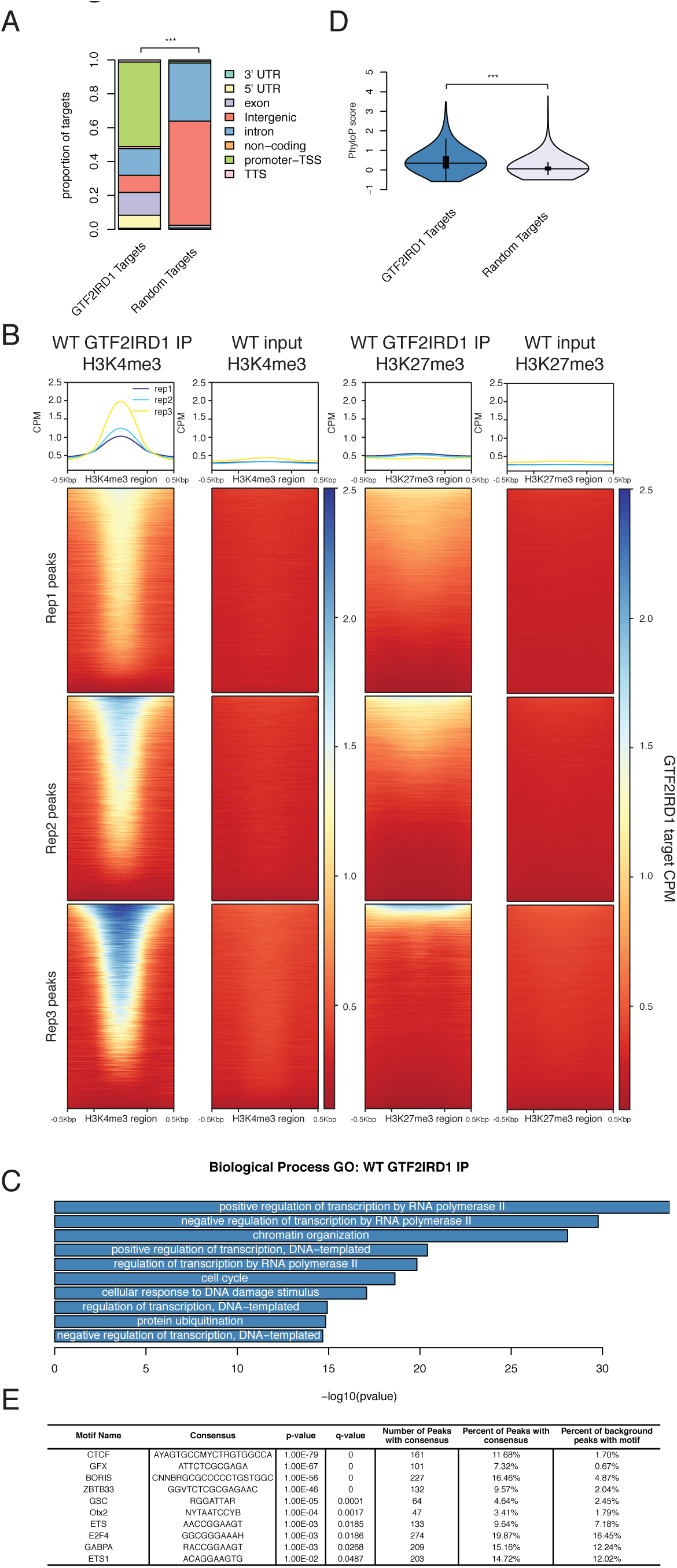
GTF2IRD1 binds preferentially to promoters in conserved, active sites in the genome. **A** GTF2IRD1 binding peaks are annotated primarily in promoters and gene bodies. The distribution of peak annotations is significantly different from random sampling of the genome. **B** GTF2IRD1 peaks were enriched in H3K4me3 sites marking active regions of the genome and to a lesser extent in H3K27me3 sites marking repressed regions. **C** GO analysis of genes that have GTF2IRD1 bound to the promoter. **D** The conservation of sequence in GTF2IRD1-bound peaks is significantly higher than expected by chance. **E** Motifs of transcription factors enriched under Gtf2ird1 bound peaks.

To understand the common functions of the genes that have GTF2IRD1 bound at the promoter we performed a GO analysis. The top ten results were consistent with the functions previously described for GTF2IRD1, specifically regulation of transcription and chromatin organization, but also identified new categories, such as protein ubiquitination (Figure 1C). To further test these regions for functional consequences, we compared the conservation of GTF2IRD1-bound peaks to a random sample of the genome and found the GTF2IRD1 peaks are more conserved (t=18.131, d.f.=2403, p < 2×10^-16^) (Figure 1D). We conducted motif enrichment analysis using HOMER to identify other factors that share binding sites with GTF2IRD1 (Figure 1E). The GSC motif, which is similar to the core RGATTR motif for GTF2I and GTF2IRD1, was identified in 4.64% of the targets (24). Interestingly, the CTCF motif was found at 11% of the GTF2IRD1 targets. These findings suggest the *Gtf2i* may modulate chromatin organization both through its direct binding at CTCF sites and through regulation of chromatin modifier genes.

GTF2I ChIP-seq showed similar results to those of GTF2IRD1. We identified 1,755 (**Supplemental Table 2**) WT GTF2I peaks that had significantly higher coverage in the WT IP compared to the KO IP (**Supplemental Figure 1A**). These peaks were significantly enriched for promoter regions as well as the gene body when compared to random genomic targets (Figure 2A)(χ^2^ = 911.63, d.f.=7, p < 2.2×10^-16^). Like GTF2IRD1, the majority of the GTF2I peaks (78.7%) overlapped H3K4me3 peaks (OR=160.98, p< 2.2×10^-16^ FET), with a smaller subset of peaks (20.7%) overlapping with the H3K27me3 mark (OR 7.022, p<2.2×10^-16^ FET). This suggests that these peaks are located mainly in active regions of the genome (Figure 2B). Summarizing the common functions of these target genes by GO analysis showed enrichment for biological processes such as intracellular signal transduction and phosphorylation (Figure 2C). For example, GTF2I binds within the gene body of the Src gene (Figure 2D), which has been shown to phosphorylate GTF2I in order to activate its transcriptional activity as well as regulate calcium entry into the cell (16, 17). The GTF2I binding sites are also significantly more conserved than random sampling of the genome, further suggesting important functional roles of these regions (Figure 2E). Motif enrichment of the GTF2I peaks revealed GC rich binding motifs, such as for the KLF and SP families of transcription factors, as well as Lhx family members. Finally, we see an enrichment of the CTCF motif, which is fitting as GTF2I has been shown to help target CTCF to specific genomic regions (25) (Figure 2F).

**Figure 2:**
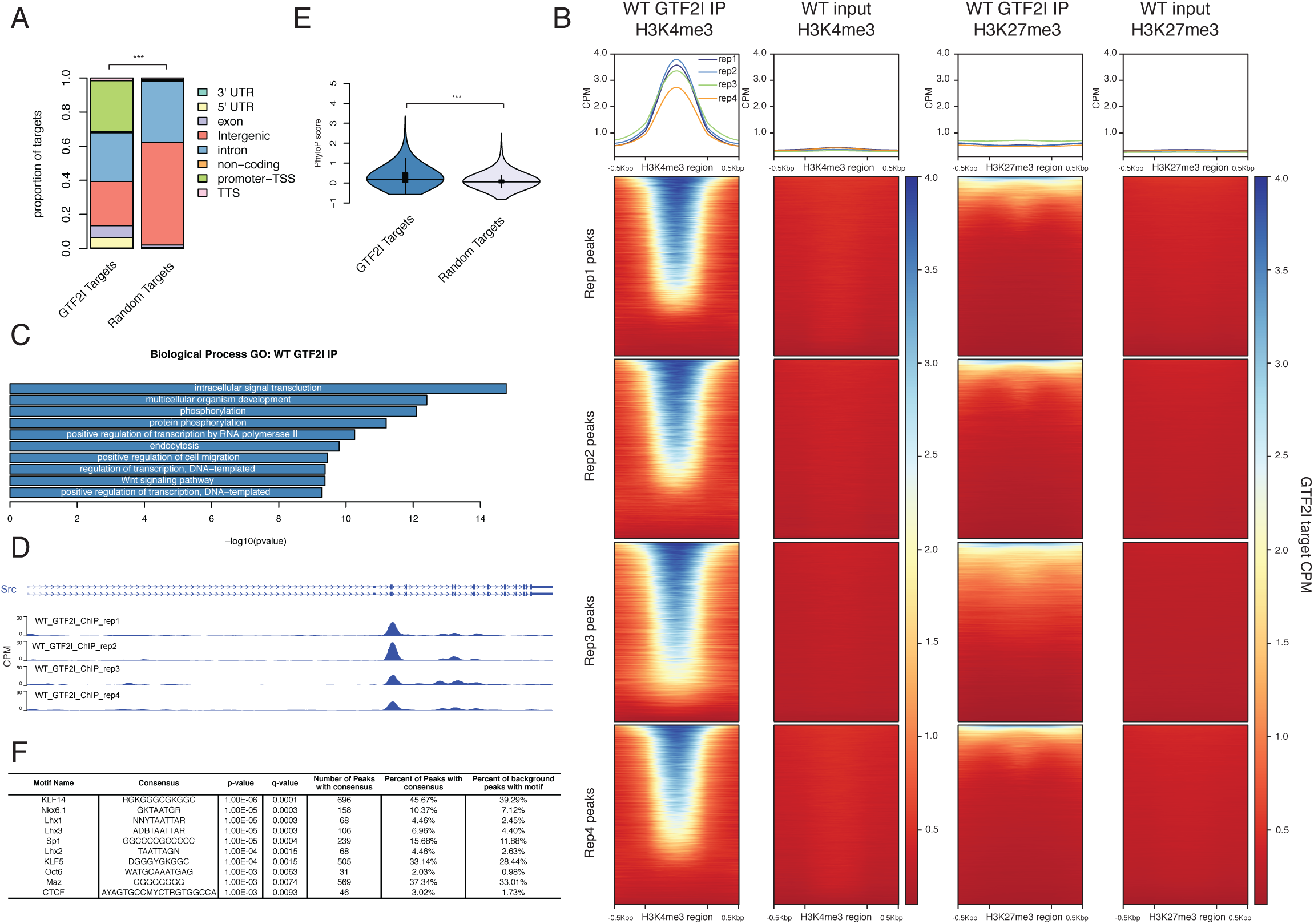
GTF2I binds at promoters in conserved, active sites in the genome. **A** GTF2I binding sites are annotated mostly in gene promoters and the gene body. The distribution of peaks is significantly different than would be expected by chance. **B** GTF2I peaks overlap with H3K4me3 peaks marking active regions and to a lesser extent GTF2I peaks fall within H3K27me3 peaks marking inactive regions. **C** GO analysis of genes that have GTF2I bound at the promoter. **D** Epigenome browser shot of GTF2I peak bound within the *Src* gene. **E** Genomic sequence under GTF2I peaks are more conserved than we would expect by chance. **F** Motifs of transcription factors that are enriched in GTF2I bound sequences.

### *Gtf2i* and *Gtf2ird1* binding sites have distinct features, yet overlap at a subset of promoters

One way in which GTF2I and GTF2IRD1 can interact is by binding the same sites in the genome. We therefore directly compared the regions bound by these proteins. First, we compared the GTF2I and GTF2IRD1 ChIP peaks and found the pattern of their binding sites is significantly different (χ^2^ = 282.84, d.f.=7, p < 2.2×10^-16^) (Figure 3A); while both transcription factors mainly bind in promoters and the gene body, GTF2IRD1 has a higher proportion of peaks at the promoter compared to GTF2I, whereas GTF2I has more peaks at intergenic regions. Interestingly, when we compared them directly to each other, the GTF2IRD1-bound peaks were significantly more conserved than the GTF2I-bound peaks (t=7.81, d.f.=2736.5, p=8.2×10^-15^) (Figure 3B). Next, to identify common targets, we identified the genes with both transcription factors at their promoter and found a significant overlap of 148 genes (p < 1×10^-38^ FET) (Figure 3C). Motif analysis on the shared peaks showed further enrichment of both CTCF and GSC motifs (Figure 3D).

**Figure 3:**
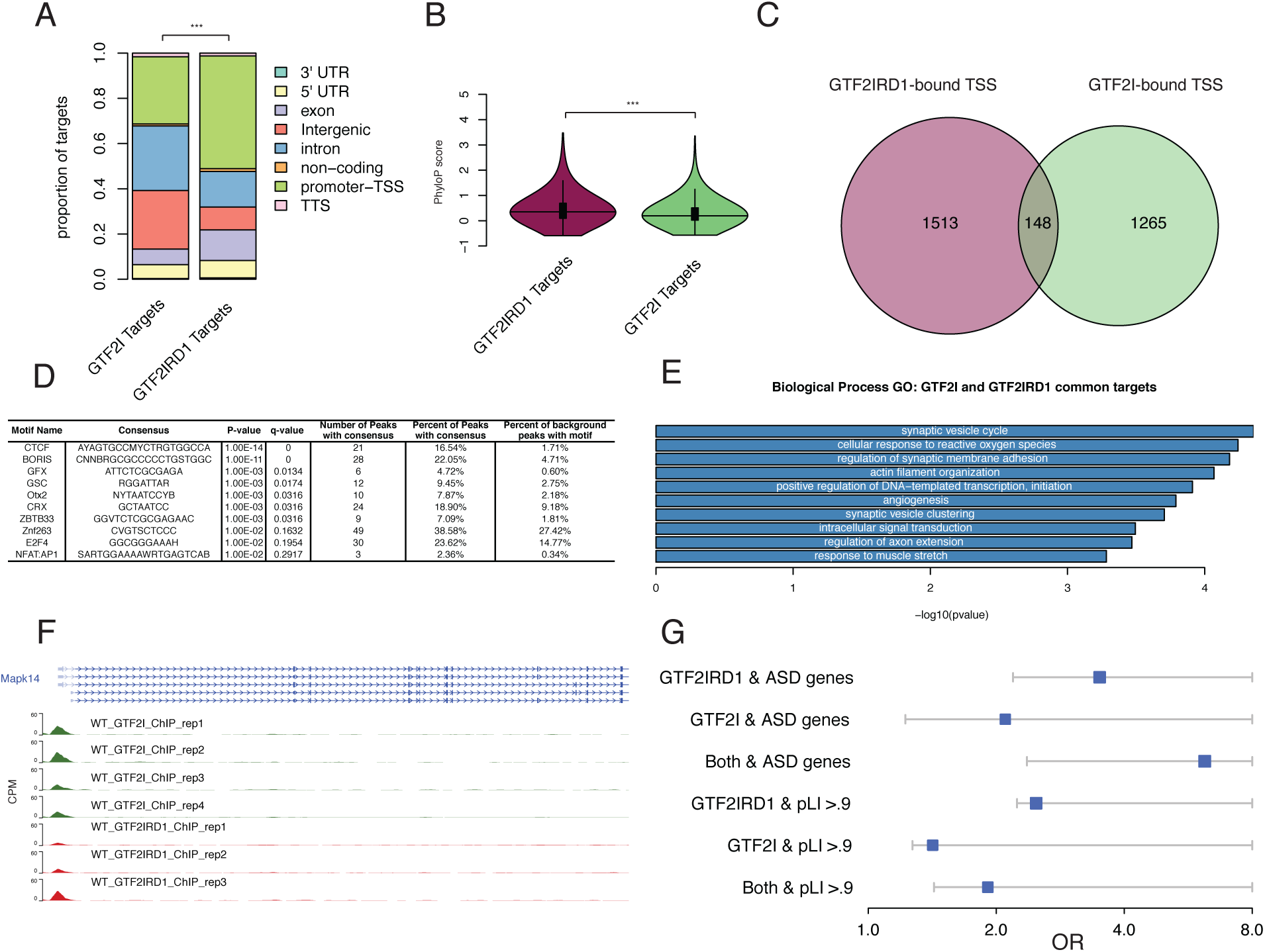
Comparison of GTF2IRD1 and GTF2I binding sites. **A** GTF2I and GTF2IRD1 have different distributions of annotated binding sites. **B** GTF2IRD1 bound sequences are more conserved than GTF2I bound sequences. **C** The overlap of genes that have GTF2I and GTF2IRD1 bound at their promoters. **D** Motifs of transcription factors that are enriched in regions bound by both GTF2I and GTF2IRD1. **E** GO analysis of genes with both GTF2I and GTF2IRD1 bound at their promoters. **F** Epigenome browser shot of *Mapk14* showing peaks for both GTF2I and GTF2IRD1. G Enrichment of GTF2IRD1 and GTF2I bound genes in ASD and conserved gene sets.

The GO functions of the overlapped genes highlight specific roles in synaptic functioning and signal transduction (Figure 3E). *Mapk14* is an example of a gene involved in signal transduction that has both GTF2I and GTF2IRD1 bound at its promoter (Figure 3F). Shared targets such as this suggest that there are points of convergence where deleting both genes, such as in WS, might result in synergistic downstream impacts.

Finally, given the consistent enrichment of CTCF binding sites in both GTF2I and GTF2IRD1 bound regions, we also compared the targets for each transcription factor to CTCF targets in E14.5 whole brain (26). We found a highly significant overlap between GTF2IRD1 and CTCF peaks, with roughly two-thirds of GTF2IRD1 binding overlapping with CTCF bound sites (939 shared peaks, OR=89.60, p < 2.2×10^-16^ FET). Similarly, GTF2I shared more peaks in common with CTCF (43%) than we would expect by chance (756 shared peaks, OR=28.16, p < 2.2×10^-16^ FET). Next, since CTCF has been shown to be present at topological associating domain (TAD) boundaries, we compared the GTF2IRD1 and GTF2I peaks with TAD boundaries determined in E14.5 cortical neurons and found significant overlaps (557 shared peaks, OR =0.16, p < 2.2×10^-16^ FET and 451 shared peaks, OR=5.19, p < 2.2×10^-16^ FET, respectively).

Thus, this TF family is remarkably enriched at promoters of neural genes, overlaps substantially at regions binding the chromatin looping protein CTCF, and is highly enriched a subset of TAD boundaries detected in the developing mouse brain. Overall, these TFs are poised to be important regulators of neural development and thus might regulate other genes associated with developmental diseases.

### *Gtf2i* and *Gtf2ird1* bind promoters of known autism genes

Copy number variations in the WS locus are clearly associated with autism spectrum disorder (27) and recent exome and genome sequencing analyses have identified over 100 more associated loci, mostly containing single genes where loss of function mutations can cause ASD (28). Interestingly, these genes are substantially enriched in chromatin modifiers and transcriptional regulators, just like GTF2IRD1 targets. However, it is unclear how mutations in this wide variety of genes all lead to a common cognitive phenotype. One widely proposed possibility is these genes are part of a functional network and mutation of any one gene might disrupt some core facet of transcriptional regulation, though most of the evidence to support this idea comes from analyses of co-expression across development (29), rather than any measure of functional interaction or targeted regulation. Therefore, we next examined whether GTF2IRD1 and GTF2I are positioned to regulate other ASD genes by binding within them or nearby. We found that GTF2IRD1 targets were highly enriched for ASD genes (OR >3.5, p-value <1.5×10^-5^ FET) – targeting 19 of the 102 genes, including a variety of epigenetic regulators such as DNMT3A, SETD5, NSD1, ADNP and SIN3A (Figure 3G). GTF2IRD1 was generally bound to their promoters or nearby CpG islands, suggesting direct regulation of these genes. GTF2I was likewise enriched at ASD genes, albeit more moderately (OR=2.1, p<.02 FET), targeting 14 genes. Five genes were targeted by both (Figure 3G). Thus our DNA binding data supports prior coexpression data that suggests a subset of ASD genes may be part of an interrelated module of chromatin modifying genes, and is consistent with prior suggestions that this convergence indicates these genes may have similar downstream pathways leading to disease (30).

It has also been found that genes mutated in ASD tend to be highly constrained in the general human population, with almost no loss of function mutations observed across >140,000 exome sequencing samples (summarized as a pLI score of >.9)(31). Strong constraint, even of heterozygous loss of function mutations, suggests that these genes do not tolerate decreased expression and thus require a very precise amount of transcription for normal human survival or reproductive fitness. We find >30% of GTF2IRD1-bound genes meet this criterion, a result highly unlikely to be due to chance (OR=2.48, p-value<2.2×10^-16^ FET). GTF2I targets also show a significant, but more modest, enrichment (OR=1.27, p-value<1.1×10^-7^ FET) (Figure 3G). Together, these results show that these TFs, especially GTF2IRD1, tend to bind genes requiring tight regulation of expression, as indicated by severe phenotypes and intolerance to loss of function mutations.

### Frameshift mutation in *Gtf2ird1* results in truncated protein and affects DNA binding at the Gtf2ird1 promoter

To investigate the functional role of *Gtf2ird1* and *Gtf2i* at these bound sites and understand how these genes interact, we made loss of function models of *Gtf2ird1* individually and a double mutant with mutations in both *Gtf2i* and *Gtf2ird1*. We designed one gRNA for each gene and injected them simultaneously into FVB/NJ mouse embryos to obtain single and double gene mutations. We first characterized the consequences of a one base pair adenine insertion in exon three of *Gtf2ird1*. This frameshift mutation introduces a premature stop codon in exon three, an early constitutively expressed exon, which we expected to trigger nonsense-mediated decay (Figure 4A). We crossed heterozygous mutant animals to analyze *Gtf2i* and *Gtf2ird1* transcript and protein abundance in heterozygous and homozygous mutants compared to WT littermates (Figure 4B). The western blots and qPCR were performed using the whole brain at E13.5. As expected, the *Gtf2ird1* mutation did not affect *Gtf2i* transcript or protein levels (Figure 4C,D). Contrary to our prediction of nonsense-mediated decay, we observed a 1.74-fold increase in the *Gtf2ird1* transcript with each copy of the mutation and a 40% reduction of the protein in homozygous mutants compared to WT with no significant difference between the WT and heterozygous mutants (Figure 4E, F). This suggests the mutation did have an effect on protein abundance and disrupted the normal transcriptional regulation of the gene, with the homozygous mutant modeling protein levels that might be expected in WS.

**Figure 4:**
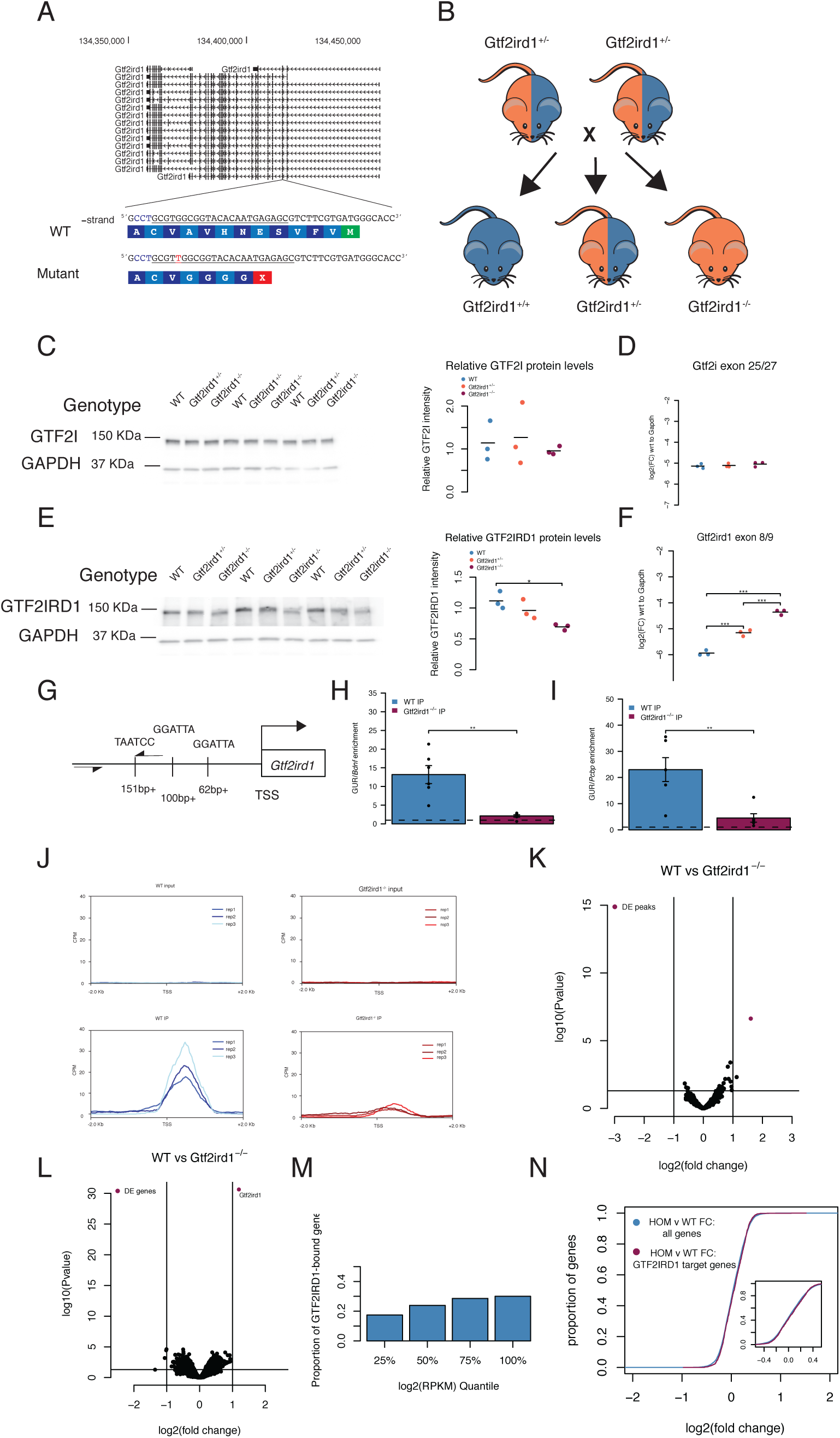
Frameshift mutation in *Gtf2ird1* exon three results in a decreased amount of an N-truncated protein with diminished binding at *Gtf2ird1* promoter and has little effect on transcription in the brain. **A** The sequence of exon three of *Gtf2ird1* targeted by the underlined gRNA with the PAM sequence in blue. The mutant allele contains a one base pair insertion of an adenine nucleotide that results in a premature stop codon. **B** Breeding scheme of the intercross of *Gtf2ird1*^+/-^ to produce genotypes used in the experiments. **C, D** Mutation in *Gtf2ird1* does not affect the protein or transcript levels of *Gtf2i*. **E** Frameshift mutation decreases the amount of protein in *Gtf2ird1*^-/-^ and causes a slight shift to lower molecular weight. **F** The abundance of *Gtf2ird1* transcript increases with increasing dose of the mutation. **G** Schematic of *Gtf2ird1* upstream regulatory element (GUR) that shows the three GTF2IRD1 binding motifs. The arrows indicate the location of the primers for amplifying the GUR in the ChIP-qPCR assay. **H,I** WT ChIP of GTF2IRD1 shows enrichment of the GUR over off target regions. There is more enrichment in the WT genotype compared to the *Gtf2ird1*^-/-^ genotype. **J** Profile plots of GTF2IRD1 ChIP-seq data confirms diminished binding at the *Gtf2ird1* promoter. **K** Differential peak analysis comparing WT and *Gtf2ird1*^-/-^ ChIP-seq data showed only the peak at *Gtf2ird1* is changed between genotypes with an FDR <0.1. **L** Differential expression analysis in the E13.5 brain comparing WT and *Gtf2ird1*^-/-^ showed only *Gtf2ird1* as changed with FDR <0.1. **M** The presence of GTF2IRD1 at gene promoters is not evenly distributed across expression levels. **N** The expression of genes bound by GTF2IRD1 is not different compared to all other genes between WT and *Gtf2ird1*^-/-^ mutants.

We noticed a slight shift in the homozygous mutant band, which may correspond to loss of the N-terminal end of the protein. Similar results were reported in another mouse model that deleted exon two of *Gtf2ird1*, in which lower levels of an N-terminally truncated protein was caused by a translation re-initiation event at methionine-65 in exon three (32). The N-terminal end codes for a conserved leucine zipper, which participates in dimerization as well as DNA-binding (32, 33). Mutating the leucine zipper was shown to affect binding of the protein to the *Gtf2ird1* upstream regulatory (GUR) element located at the promoter of *Gtf2ird1*. Given the previous findings that GTF2IRD1 negatively autoregulates its own transcription and mutating the leucine zipper affects binding to the GUR, we hypothesized that the frameshift mutation diminished the ability of GTF2IRD1 to bind its promoter resulting in increased transcript abundance. We tested this by performing ChIP-qPCR in the E13.5 brain in WT and *Gtf2ird1*^-/-^ mutants. In the WT brain, GTF2IRD1 IP enriched for the GUR 13-20 times over off-target sequences, which was significantly higher than the GTF2IRD1 IP in the *Gtf2ird1*^-/-^ brain (Figure 4G,H,I). Taken together, nonsense transcripts of *Gtf2ird1* with a stop codon in exon 3 can reinitiate at a lower level to produce a N-truncated protein with diminished binding capacity at the GUR element.

### Truncated *GTF2IRD1* does not affect binding genome wide

Given that the one base pair insertion did not result in a full knockout of the protein but did affect its DNA binding capacity at the GUR of *Gtf2ird1*, we tested whether the mutant was a loss of function for all DNA binding. We performed ChIP-seq in the E13.5 *Gtf2ird1*^-/-^ mutants and compared it to WT ChIP-seq data. This comparison confirmed the decrease in binding at the TSS of *Gtf2ird1*, suggesting the mutation has greatly decreased binding at this locus (Figure 4J). Surprisingly, the only peak identified as having differential coverage (FDR < 0.1) between the two genotypes was the peak at the *Gtf2ird1* TSS (Figure 4K). This suggests that the frameshift mutation has a very specific consequence on how GTF2IRD1 binds to its own promoter that does not robustly affect its binding elsewhere in the genome. The *Gtf2ird1* promoter has two instances of the R4 core motif in the sense direction and one instance of the motif in the antisense orientation. We searched the sequences under the identified peaks for similar orientations of this binding motif and found three other peaks, none of which showed any difference in binding coverage between genotypes. However, these three other peaks did not match the exact spacing of the R4 motifs found in the *Gtf2ird1* promoter. This suggests that the leucine zipper is important for a specific configuration of binding sites that is only present in this one instance in the genome. It has been shown that the three R4 motifs are present in the same orientation and spacing across great evolutionary distances (32).

### *Gtf2ird1* frameshift mutation shows mild transcriptional differences

The N-truncation of GTF2IRD1 clearly affected its binding at the *Gtf2ird1* promoter and affected expression levels. Although we didn’t see genome-wide binding perturbed, it is possible losing the N terminus, or the decreased protein level, altered the protein’s ability to recruit other transcriptional co-regulators to impact gene expression. Therefore, we tested the effects of this mutation on genome-wide transcription in the E13.5 brain. We compared the whole brain transcriptome of WT littermates to heterozygous and homozygous mutants. Strikingly similar to ChIP-seq data, the only transcript with an FDR < 0.1 was *Gtf2ird1*, which was affected in the same direction seen in the qPCR results (Figure 4L and **Supplemental Figure 2A**). We leveraged WT ChIP-seq data to test if GTF2IRD1 presence at a promoter correlates with gene expression. Binning the genes according to expression level showed that the distribution of GTF2IRD1 targets was different than expected by chance (χ^2^ = 48.83, d.f.=3 p < 1.42×10^-10^), suggesting highly expressed genes are more likely to have GTF2IRD1 bound at their promoters (Figure 4M). The majority (1262 peaks) of the GTF2IRD1 peaks at a TSS were at expressed genes, with only 410 next to genes not expressed at detectable levels in the E13.5 brain. To see if there was a more subtle general effect below our sensitivity to detect by analysis of single genes, we tested the bound GTF2IRD1 targets expressed as a population for a shift in expression. We saw a trend towards significance between the bound and unbound genes, but with a small effect size: a mean increase of 0.014 fold change in GTF2IRD1 targets (Kolgmogorov-Smirnov test D=0.038, p=0.079). While this is perhaps unsurprising because the frameshift mutation did not disturb binding genome wide (Figure 4N), the homozygous mutants do have an overall decrease in protein level of ∼50%, which should mimic a WS deletion. Thus, transcriptional consequences of hemizygosity of this gene might be similarly small.

### Frameshift mutation is sufficient to affect behavior

Although we observed only small differences in DNA binding and overall brain transcription, another *Gtf2ird1* model also reported little to no effects of mutation on transcriptome-wide expression in the brain, yet the model still showed behavioral phenotypes (8, 34). Therefore, we tested our mutation for downstream consequences on adult mouse behavior (Table 1). There are many single gene knockout models of *Gtf2ird1* and each shows distinct behavioral differences which are, in some instances, contradictory (8,9,21,35). One consistent phenotype across models is a deficit in motor coordination, which is also affected in individuals with WS. Similarly, we observed a significant effect of genotype (H_2_=16.35, p=0.0003), on balance. Heterozygous and homozygous animals fell off a ledge sooner than WT littermates (p=0.0038, p=0.0007, respectively) (Figure 5A). Marble burying has not been reported in other *Gtf2ird1* models, but in larger WS models that delete the entire syntenic WSCR or the proximal half of the region containing *Gtf2ird1* have shown decreased marble burying in mutants (5, 36). We observed a similar significant effect of genotype on the number of marbles buried (F_2,80_=6.17, p =0.0033), with *Gtf2ird1*^-/-^ mutants burying fewer marbles than WT mice (p=0.002) (Figure 5B). Reports of overall activity levels in *Gtf2ird1* mouse models have been discrepant (9, 21). Here we show there is no main effect of genotype (F_2,88_=1.36, p=0.263) but a time by genotype interaction (F_10,440_=5.791, p=3.3×10^-8^) on total distance traveled in a one-hour locomotor task. Though no single time point showed a difference between genotypes when performing post hoc tests, this may reflect an overall difference in habituation to the chamber (Figure 5C). When taking sex into consideration, there was a main effect of sex (F_1,85_=5.23, p=0.025) and the genotype by time interaction persists (F_10,425_=5.82, p=3.06×10^-8^) (**Supplemental Figure 3 A,B**). Time spent in the center of an open field is used as a measure of anxiety-like behavior in mice. Anxiety-like behaviors in *Gtf2ird1* models have also been discrepant in the literature (8). Here we show that there was no main effect of genotype on center variables in this task (F_2,88_=0.88, p=0.42) (Figure 5D).

**Table 1:** Behaviors and sample sizes for *Gtf2ird1*^+/-^ x *Gtf2ird1*^+/-^ cross

| Cohort |  | male |  |  | female |  |  |
| --- | --- | --- | --- | --- | --- | --- | --- |
| Behavior | Experimenter | WT | *Gtf2ird1*<sup>+/-</sup> | *Gtf2ird1*<sup>-/-</sup> | WT | *Gtf2ird1*<sup>+/-</sup> | *Gtf2ird1*<sup>-/-</sup> |
| 1-hour locomotor activity | Female | 10 | 21 | 12 | 15 | 16 | 17 |
| Ledge | Female | 9 | 21 | 13 | 15 | 17 | 17 |
| Marble burying | Female | 8 | 20 | 10 | 14 | 17 | 14 |
| Pre-pulse inhibition | Female | 8 | 20 | 11 | 14 | 17 | 15 |
| Conditioned fear | Female | 9 | 18 | 10 | 15 | 17 | 17 |
| Shock sensitivity | Female | 9 | 10 | 11 | 15 | 17 | 17 |

**Figure 5:**
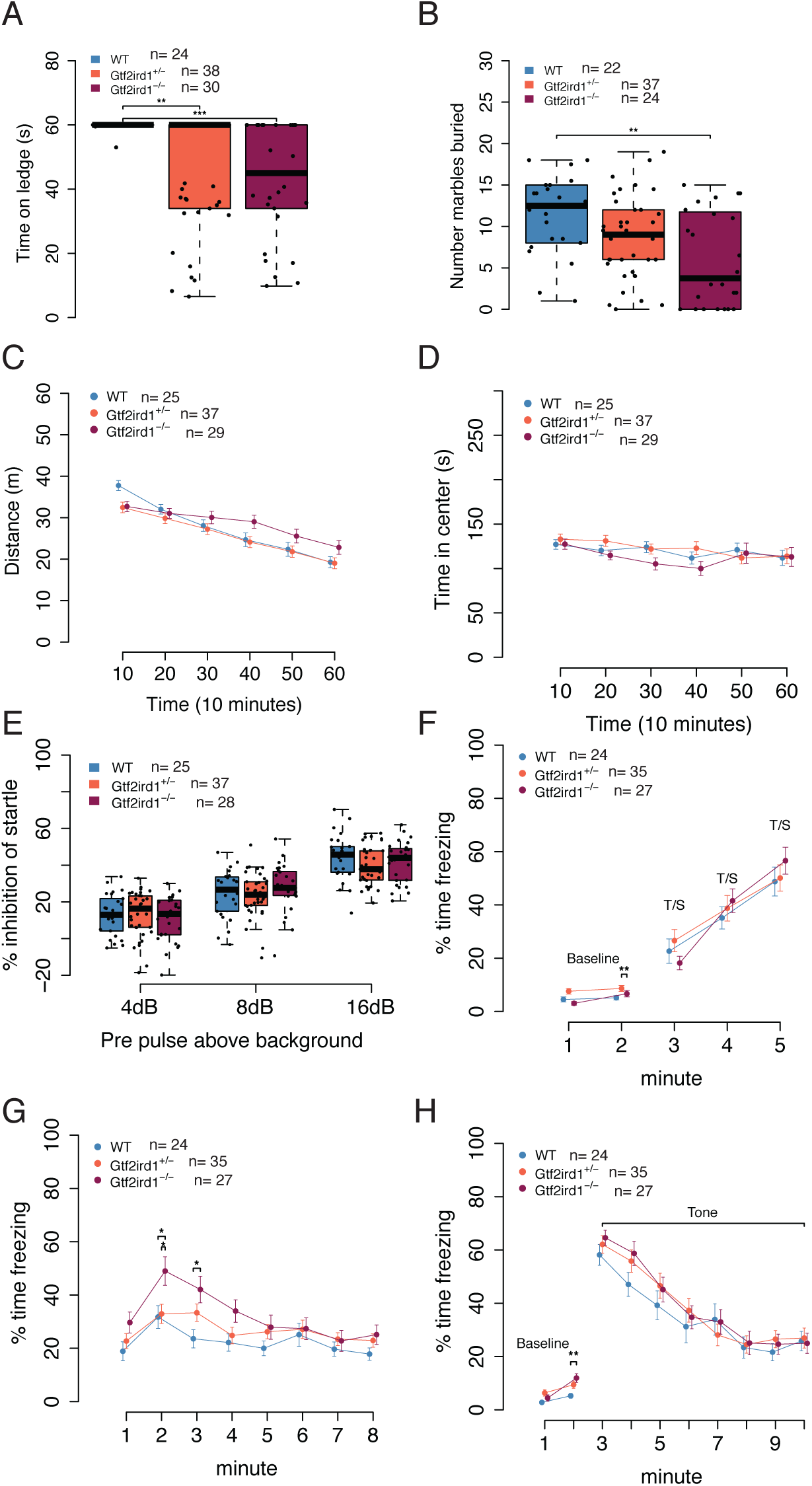
Homozygous frameshift mutation in *Gtf2ird1* is sufficient to cause behavioral phenotypes. **A** Homozygous mutants have worse balance than WT littermates in ledge task. **B** Homozygous mutants bury fewer marbles than WT and heterozygous littermates. **C** Overall activity levels are not affected but a time by genotype interaction shows the mutant animals are slower to habituate to the novel environment. **D** There is no difference in time spent in the center of the apparatus between genotypes. **E** All animals show an increase in startle inhibition when given a pre-pulse of increasing intensity. There is no difference between genotypes. **F** Acquisition phase of fear conditioning paradigm. All animals show the expected increase in freezing to additional foot shocks. **G** *Gtf2ird1*^-/-^ animals show an early increased contextual fear memory response compared to WT and heterozygous littermates. **H** There were no significant differences between genotypes in cued fear.

Finally, as individuals with WBS also show a high prevalence of phobias, sensitivity to sounds, and learning deficits (3, 37), we tested sensory motor gating and learning and memory. Pre-pulse inhibition (PPI) results from a decreased startle to an auditory stimulus when the startle stimulus is preceded by a smaller stimulus. PPI was reduced in animals with the proximal WSCR region deleted, but mice with the distal deletion or full deletion did not have any abnormal phenotype (6). In our study, there was no main effect of genotype on PPI (F_2,87_=0.24, p=0.79), but a pre-pulse by genotype interaction (F_4,174_=2.66, p=0.034), which suggests that for some pre-pulse stimuli there is a difference between mutants and WT littermates, however no comparisons survived multiple testing corrections in the post hoc test (Figure 5E).

In our assessment of learning and memory with the conditioned fear paradigm, there was a main effect of genotype (F_2,83_=4.82, p=0.010) and minute (F_1,83_=9.75, p=0.002) on baseline freezing. Post hoc tests on baseline data showed the heterozygous mutants froze more than homozygous mutants during minute one (p=0.0065). After the baseline, when animals were trained to associate a tone with a footshock, we observed all mice had increased freezing over time (F_2,122_=26.77, p=2.28×10^-10^) as expected. (Figure 5E). On the second day, which tested contextual fear memory, all genotypes exhibited a fear memory response as indicated by the significant effect of the context compared to baseline (F_1,83_=173.20, p<2×10^-16^). Each group froze more during the first two minutes of day two than on day one (WT: p=1.98×10^-8^, *Gtf2ird1*^+/-^: p=4.97×10^-11^, *Gtf2ird1*^-/-^: p<2×10^-16^) (**Supplemental Figure 3C**). When we analyzed the entire time of the experiment of contextual fear we similarly saw no main effect of genotype (F_2,83_=2.8946, p=0.061), but a significant effect of time (F_7,581_=15.05, p<2×10^-16^) and a time by genotype interaction (F_14,581_=2.01, p=0.016). Post hoc analysis showed that during minute two the homozygous animals froze significantly more than the *Gtf2ird1*^+/-^ mutants (p=0.026). Similarly, the homozygous mutants froze more than WT littermates during minutes two (p=0.021) and three (p=0.012), suggesting an increased contextual fear memory response (Figure 5G). On day three of the experiment, we tested cued fear. During day three baseline we saw a difference in freezing between genotypes (F_2,83_=4.13, p=0.02) as well as a time by genotype interaction (F_2,83_=4.47, p=0.014), with homozygous mutants freezing more in minute two than WT littermates (p=0.002). All genotypes had a similar response to the tone (F_2,83_=0.36, p=0.70) (Figure 5H). These differences could not be explained by differences in shock sensitivity (flinch: H_2_=2.52, p=0.28, escape: H_2_=3.13, p=0.21, vocalization: H_2_=2.20, p=0.33) (**Supplemental Figure 3D**). Thus, mutation of *Gtf2ird1* appears to enhance contextual fear learning.

Overall, these behavioral analyses show the N-terminal truncation and/or decreased protein levels of the *Gtf2ird1* mutant still result in adult behavioral phenotypes, specifically in the domains of balance, marble burying, and fear conditioning. The most severe phenotypes were observed in the homozygous mutants, which may model the haploinsufficiency of WS deletions.

### Generation of a *Gtf2i* and *Gtf2ird1* double mutant

The evidence of functional consequences from the one base pair *Gtf2ird1* frameshift mutation led us to characterize a double mutant that was generated during the dual gRNA CRISPR/Cas9 injections. This mutant allowed us to test the effects of knocking out *Gtf2i* combined with a *Gtf2ird1* mutation, and test different *Gtf2ird1* mutations for consistency in phenotypes. The double mutant described here has a two base pair deletion in exon five of *Gtf2i* and a 589bp deletion that encompasses most of exon three of *Gtf2ird1* (Figure 6A). We carried out a heterozygous cross of the double mutants to test the protein and transcript abundance of each gene in the heterozygous and homozygous states. The homozygous double mutant is embryonic lethal due to the lack of *Gtf2i*, which has been described in other *Gtf2i* mutants (Figure 6B) (4, 38). We were, however, able to detect homozygous embryos up to E15.5. Thus, we focused molecular analyses on E13.5 brains. The two base pair deletion in exon five of *Gtf2i* leads to a premature stop codon resulting in a full protein knockout, and decreases the transcript abundance consistent with degradation of the mRNA due to nonsense-mediated decay (Figure 6C,D). The 589bp deletion in *Gtf2ird1* removes all of exon three except the first 14bp. We observed the same increase in transcript abundance that was detected in the one base pair insertion mutation but this mutation had a larger effect on protein levels across genotype, with homozygous mutants producing a truncated protein at about 10% of WT levels (Figure 6E,F).

**Figure 6:**
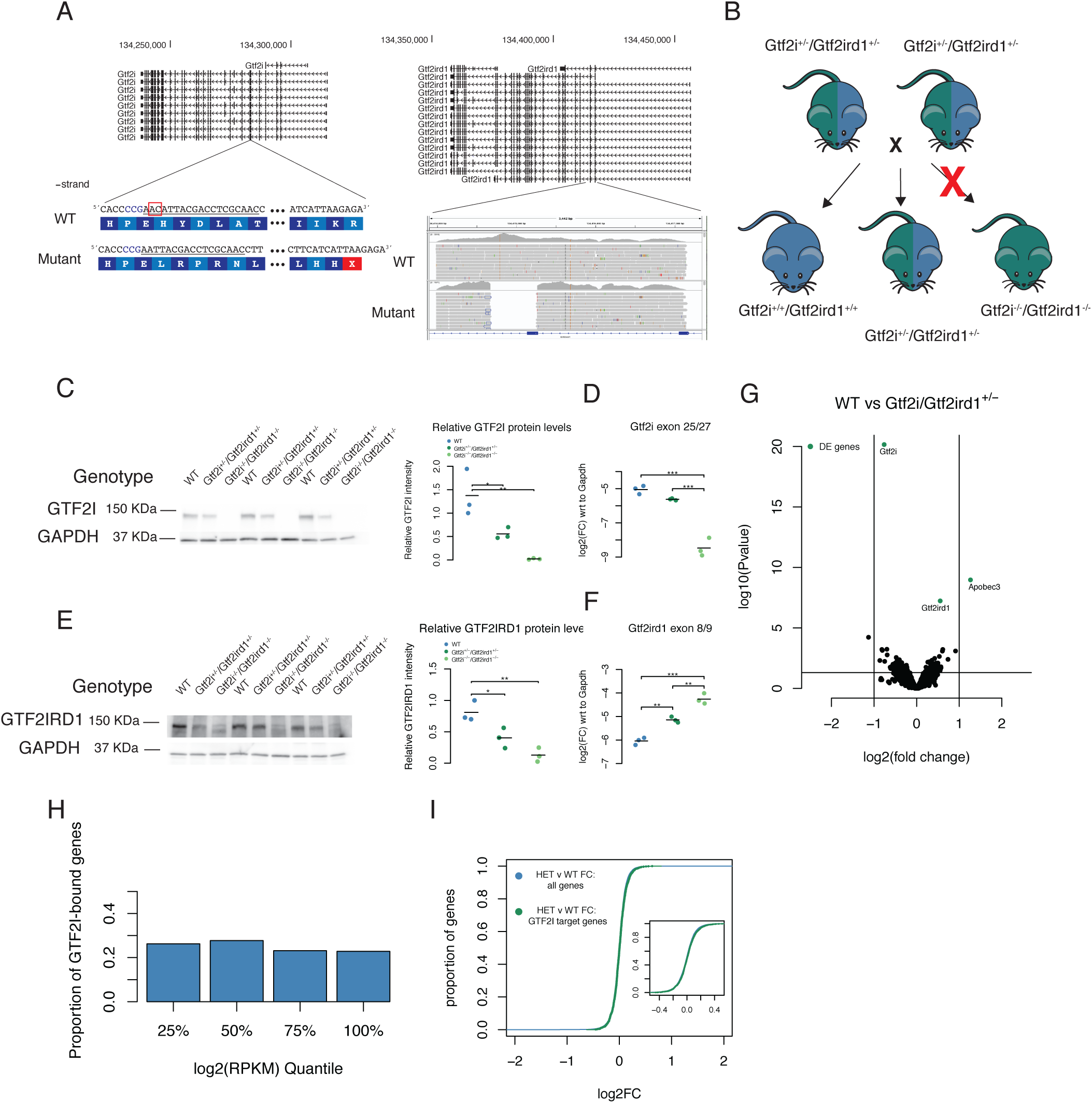
Mutating both *Gtf2i* and *Gtf2ird1* does not result in larger differences in brain transcriptomes. **A** Generation of double mutant. gRNA target is underlined in exon five of *Gtf2i* with the PAM sequence in blue. The two base pair deletion results in a premature stop codon within exon five. The *Gtf2ird1* mutation is a large 589bp deletion covering most of exon three as shown in the IGV browser shot. **B** Heterozygous intercross to generate genotypes for ChIP and RNA-seq experiments. The homozygous double mutants are embryonic lethal but are present up to E15.5. **C** The two base pair deletion in *Gtf2i* decreases the protein by 50% in heterozygous mutants and no protein is detected in the homozygous E13.5 brain. **D** The mutation decreases the abundance of *Gtf2i* transcript consistent with nonsense-mediated decay. **E** The 589bp deletion in *Gtf2ird1* leads to decreased protein levels in heterozygous and homozygous mutants. There is still a small amount of protein made in the homozygous mutant. **F** The 589bp deletion increases the amount of *Gtf2ird1* transcript. **G** Volcano plot comparing the expression in the E13.5 brain of WT and heterozygous double mutants. The highlighted genes represent an FDR <0.1. **H** The presence of *Gtf2i* at the promoters does not correlate with the expression of a gene. **I** The fold change of genes between WT and heterozygous double mutants that have GTF2I bound at their promoters were slightly upregulated when compared to the fold change of genes that did not have GTF2I bound.

### Knocking down both *Gtf2i* and *Gtf2ird1* produces mild transcriptomic changes

To test if combined mutation of *Gtf2i* and *Gtf2ird1* had a larger effect on the transcriptome, we performed whole brain RNA-seq analysis on WT E13.5 brains and compared them to *Gtf2i*^+/-^/*Gtf2ird1*^+/-^ littermates. Similar to what was seen in the previous *Gtf2ird1*^-/-^ mutants, there were only mild differences between the transcriptomes (Figure 6G). We also compared WT transcriptomes to the homozygous double mutants, which showed greater differences. However, since these mutants have a very severe phenotype, including neural tube closure defects, any direct transcriptional consequences are probably masked by a large number of indirect effects. Indeed, GO term analysis suggested that overall nervous system development and glial cell differentiation is disrupted (**Supplemental Figure 4A,B**). We also analyzed GTF2I ChIP-seq data with RNA-seq data. Unlike the enriched binding at highly expressed genes we saw with GTF2IRD1 alone, gene expression levels were not significantly related to GTF2I binding (χ^2^ = 6.58, d.f.=3 p=0.086) (Figure 6H). This is consistent with a previous report of GTF2I ChIP-seq data. Again, the majority (963 peaks) of the TSS GTF2I peaks were nearby expressed genes, with 458 next to genes not expressed at detectable levels in the E13.5 brain. There is a slight but significant increase in gene expression of genes bound by GTF2I compared to genes that are not (0.0213 logFC, Kolgmogorov-Smirnov test D=0.075, p=9.50×10^-5^) (Figure 6I). Thus, heterozygous *Gtf2i* and *Gtf2ird1* mutation, like Gt2ird1 mutation alone, results in very subtle transcriptional changes.

### Double mutants show behavioral consequences similar to single *Gtf2ird1* mutants

To test the effects of mutating both *Gtf2i* and *Gtf2ird1* on behavior, we crossed the heterozygous double mutant to the single *Gtf2ird1* heterozygous mouse (Figure 7A). This breeding strategy produced four littermate genotypes, WT, *Gtf2ird1*^+/-^, *Gtf2i*^+/-^/*Gtf2ird1*^+/-^, and *Gtf2i*^+/-^/*Gtf2ird1*^-/-^ for direct and well-controlled comparisons. To test for additive effects, the primary comparison was contrasting *Gtf2ird1*^+/-^ mice to *Gtf2i*^+/-^/*Gtf2ird1*^+/-^ littermates. The remaining genotype, *Gtf2i*^+/-^ /*Gtf2ird1*^-/-^, further tested the effects of heterozygous *Gtf2i* mutation in the presence of both *Gtf2ird1* mutations. To be thorough, we tested protein and transcript abundance of each gene in all four genotypes. As expected, all genotypes with the *Gtf2i* mutation showed decreased protein and transcript levels. The *Gtf2ird1* results reflected what was previously shown for each mutation, however, *Gtf2i*^+/-^/*Gtf2ird1*^-/-^ did not show any further detectable decrease in protein abundance compared to the *Gtf2i*^+/-^/*Gtf2ird1*^+/-^ genotype (**Supplemental Figure 5A-D**), with both at about 50% of WT levels.

**Figure 7:**
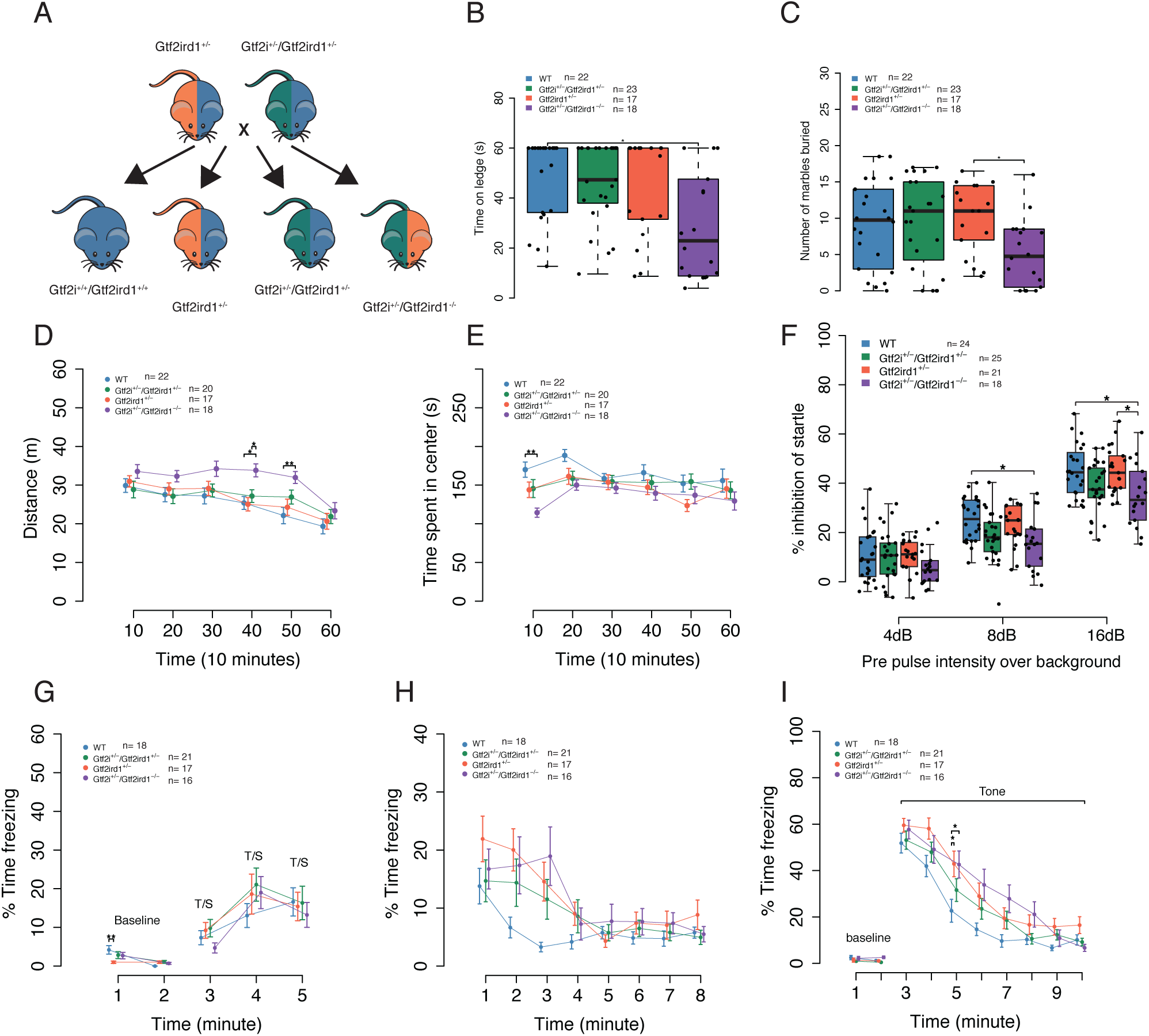
*Gtf2i* does not modify most of the phenotypes of *Gtf2ird1* mutation. **A** Breeding scheme for behavior experiments. **B** The *Gtf2i*^+/-^/*Gtf2ird1*^-/-^ animals fell off ledge sooner than WT littermates. **C** There was a main effect of genotype on marbles buried. Post hoc analysis showed that *Gtf2i*^+/-^/*Gtf2ird1*^-/-^ buried fewer marbles than the *Gtf2ird1*^-/-^ genotype. **D** *Gtf2i*^+/-^/*Gtf2ird1*^-/-^ had increased overall activity levels in a one-hour activity task. **E** *Gtf2i*^+/-^/*Gtf2ird1*^-/-^ showed decreased time in the center of the apparatus compared to WT. **F** All animals show an increased startle inhibition when given a pre-pulse of increasing intensity. The double mutants show less of an inhibition at higher pre-pulse levels compared to WT and *Gtf2ird1*^+/-^ animals. **G** All genotypes showed increased freezing with subsequent foot shocks. **H** All genotypes showed a similar contextual fear response. **I** There was a main effect of genotype on cued fear with the *Gtf2ird1*^+/-^ and *Gtf2i*^+/-^/*Gtf2ird1*^-/-^ genotypes showing an increased fear response compared to WT.

We repeated the same behaviors performed on the one base pair *Gtf2ird1* mutants (Table 2). We saw a similar significant effect of genotype on balance (H_3_=10.68, p=0.014), with *Gtf2i*^+/-^/*Gtf2ird1*^-/-^ mice falling off sooner compared to WT littermates (p=0.025) (Figure 7B). There was no significant difference between the *Gtf2ird1*^+/-^ and *Gtf2i*^+/-^/*Gtf2ird1*^+/-^ genotypes, suggesting that decreasing the dosage of GTF2I does not strongly modify the *Gtf2ird1*^+/-^ phenotype. There was a significant effect of genotype on the number of marbles buried (F_3,76_=2.93, p=0.039). Post hoc analysis showed a significant difference between only *Gtf2ird1*^+/-^ and *Gtf2i*^+/-^ /*Gtf2ird1*^-/-^ littermates (p=0.050) (Figure 7C), with a trend in the same direction as was previously seen in the *Gtf2ird1*^-/-^ mutants. We saw a main effect of genotype on activity levels in the one-hour locomotor task (F_3,69_=3.22, p=0.028), but we did not see the same main effect of sex (F_1,69_=2.29, p=0.14), or a sex by genotype interaction (F_3,69_=1.82, p=0.15); however we did see a three-way sex by time by genotype interaction (F_15,345_=1.95, p=0.018). The combined sex data showed *Gtf2i*^+/-^/*Gtf2ird1*^-/-^ mice travel a greater distance than WT and *Gtf2ird1*^+/-^ mice at time point 40 (Figure 7D). When we looked at the data by sex, we saw a larger effect in females with *Gtf2ird1*^+/-^ and *Gtf2i*^+/-^/*Gtf2ird1*^+/-^ genotypes intermediate to *Gtf2i*^+/-^/*Gtf2ird1*^-/-^ (**Supplemental Figure 5E,F**). There was also a main effect of genotype on the time spent in the center of the apparatus (F_3,69_=3.60, p=0.018) that was not seen in the previous *Gtf2ird1* cross. *Gtf2i*^+/-^/*Gtf2ird1*^-/-^ mice spent less time in the center during the first ten minutes of the task compared to WTs (p=0.0019) (Figure 7E).

**Table 2:**
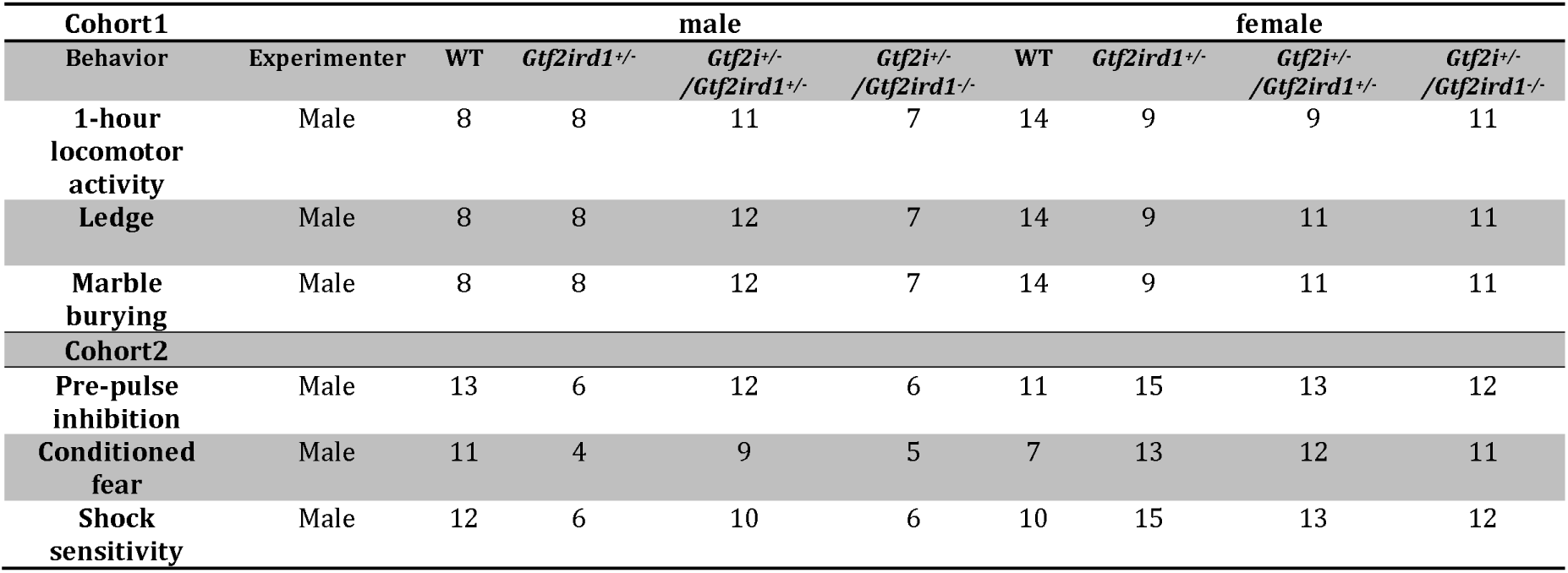
Behaviors and sample sizes for *Gtf2ird1*^+/-^ x *Gtf2i*^+/-^/*Gtf2ird1*^+/-^ cross

Finally, we repeated the PPI and conditioned fear memory tasks using this breeding strategy. In contrast to what was observed for PPI in the *Gtf2ird1* cross, we saw a significant main effect of genotype (F_3,84_=4.59, p=0.0051). The *Gtf2i*^+/-^ /*Gtf2ird1*^-/-^ mice showed an attenuated PPI response especially at the louder pre-pulse stimuli compared to WT littermates (PPI8:p=0.02, PPI16:p=0.018) and *Gtf2ird1*^+/-^ mice (PPI16:p=0.02) (Figure 7F). On day one of the conditioned fear task, all genotypes showed increased freezing with subsequent foot shocks as expected. WT animals exhibited higher freezing during minute one of baseline, but this difference diminished during minute two (Figure 7G). All animals showed a contextual fear memory response when they were re-introduced to the chamber on day two (F_1,68_=81.21, p=3.21×10^-13^) (**Supplemental Figure 5G**). While there was no main effect of genotype (F_3,68_=1.61, p=0.19) (Figure 7H), the *Gtf2ird1*^+/-^ and double mutants showed a trend towards increased freezing that was seen in the previous behavior cohort. On day three, when cued fear was tested, there was a significant effect of genotype on the freezing behavior (F_3,68_=3.17, p=0.030) and a time by genotype interaction (F_21,476_=1.63, p=0.040). During minute five of the task the *Gtf2i*^+/-^/*Gtf2ird1*^-/-^ mutants froze significantly more than WTs (p=0.030) as did the *Gtf2ird1*^+/-^ (p=0.024) (Figure 7I). The cued fear phenotype could not be explained by differences in shock sensitivity (**Supplemental Figure 5H**).

By crossing these mutant lines, we tested the hypothesis that the double heterozygous mutant would be more severe than a mutation only affecting *Gtf2ird1*. *Gtf2ird1*^+/-^ and *Gtf2i*^+/-^/*Gtf2ird1*^+/-^ genotypes resulted in mild deficits compared to WTs that, in some cases, were intermediate to the *Gtf2i*^+/-^/*Gtf2ird1*^-/-^ phenotype. There were no instances where the *Gtf2ird1*^+/-^ or *Gtf2i*^+/-^/*Gtf2ird1*^+/-^ genotypes were significantly different from each other, suggesting that in the behaviors we have tested, the *Gtf2i* mutation does not modify the effects of a *Gtf2ird1* mutation. This unique cross also allowed us to characterize a new mouse line *Gtf2i*^+/-^/*Gtf2ird1*^-/-^, which had the largest impact on behaviors. The phenotypes of *Gtf2i*^+/-^/*Gtf2ird1*^-/-^ were always in the same direction as the phenotypes in the *Gtf2ird1*^-/-^ mouse model, but we also saw a significant PPI deficit and cued fear difference when the *Gtf2i* mutation was added. This further supports that the behaviors tested here, such as, balance, marble burying, and learning and memory are largely affected by homozygous mutations in *Gtf2ird1*.

## Discussion

We have described the in vivo DNA binding sites of GTF2IRD1 and GTF2I in the developing mouse brain. This is the first description of these two transcription factors in a tissue that is relevant for the behavioral phenotypes seen in mouse models of WS. GTF2IRD1 showed a preference for active sites and promoter regions. The conservation of GTF2IRD1 targets was higher on average than would be expected by chance, which provides evidence that these are functionally important regions of the genome. The functions of genes bound by GTF2IRD1 include transcriptional regulation, such as chromatin modifiers, as well as post-translational regulation including protein ubiquitination. A role for GTF2IRD1 in regulating genes involved in protein ubiquitination has not been described before. This supports the role of GTF2IRD1 in regulating chromatin by transcriptionally controlling other chromatin modifiers. These data, along with the localization pattern of GTF2IRD1 in the nucleus and its direct association with other chromatin modifiers such as ZMYM5 (18, 39), suggests that GTF2IRD1 can exert its regulation of chromatin at several different levels of biological organization. Motif enrichment analysis of GTF2IRD1 peaks indicated that CTCF cobind with GTF2IRD1. Consistent with this, GTF2IRD1 is also often present at TAD boundaries where CTCF is known to be enriched. This further suggests GTF2IRD1 may have a role in defingin chromatin topology. GTF2I has been shown to interact with and target CTCF to specific sites in the genome (25) so it would be interesting to test if GTF2IRD1 has a similar relationship with CTCF.

Overall, GTF2I showed a similar preference for promoters and active regions, although it had more intergenic targets than GTF2IRD1, and the conservation of GTF2I peaks was significantly lower than GTF2IRD1 peaks. The genes bound by GTF2I were enriched for signal transduction and phosphorylation. Interestingly, GTF2I was bound to the Src gene body. SRC is known to phosphorylate GTF2I to induce its transcriptional activity (16). Phosphorylation of GTF2I by SRC also antagonizes calcium entry into the cell (17). While knocking out *Gtf2i* did not affect the expression of Src, it would be interesting to understand the functional consequence of GTF2I binding Src, especially since Src knockout mice exhibit similar behaviors as *Gtf2i* mouse models (40).

The overlap of GTF2I and GTF2IRD1 targets was significant, and the target genes were enriched for synaptic activity and signal transduction. This is evidence that these genes can interact via their binding targets to produce cognitive and behavioral phenotypes. To test the difference between combined *Gtf2i* and *Gtf2ird1* mutation and mutation of *Gtf2ird1* alone, we characterized two new mouse models. We used the CRISPR/Cas9 system to generate multiple mutations in the two genes individually as well as together from one embryo injection. The ease and combinatorial possibilities of this technology will be amenable to testing many unique combinations of genetic mutations in copy number variant regions, which will be important to fully understand the complex relationships of genes in these disorders.

We found a frameshift mutation expected to trigger non-sense mediated decay in *Gtf2ird1* did not degrade the mRNA but did result in an N-terminal truncation and protein level reduction in the homozygous mutant (23). Even a larger, 589bp deletion of exon three in *Gtf2ird1* did not result in mRNA degradation, but did have a larger effect on protein level. This phenomenon of increased *Gtf2ird1* RNA levels has been seen in at least three other mouse models of *Gtf2ird1* (8,23,32). Two of these were made using classic homologous recombination removing either exon two alone or exon two through part of exon five. In both of these models, *Gtf2ird1* transcript was still made, but no in vivo protein analysis was done due to poor quality antibodies and the undetectably low protein expression in WT mice. The third model also saw the N-terminal truncation. The presence of an aberrant protein that can still bind the genome, such as the mutant described here, could explain the lack of transcriptomic differences in the brain shown here and by others (34). The mutant protein may also still interact with other binding partners and be trafficked to the appropriate genomic loci. This mutation did disrupt the binding of GTF2IRD1 to its own promoter, which resulted in an increase in transcript levels. The property that specifies GTF2IRD1 binding to its own promoter must be unique, as DNA binding genome-wide was not robustly perturbed in the mutant.

In the end, GTF2IRD1 has proven to be a remarkably difficult protein to disrupt in a targeted manner – a finding that may modify interpretation of prior studies using a variety of mutant lines, including ours (23). Indeed, it took years to establish a sufficiently sensitive immunoblotting protocol for GTF2IRD1, much less a ChIP-seq protocol to study its binding genome wide. Thus, even in the current study, much of the transcriptional and behavioral characterization was complete prior to discovering that substantial DNA binding remained. Presumably the resiliency of this binding also extends to the mutants we used in our recent test of the sufficiency of *Gtf2i* family mutants to recapitulate the deletion of the entire locus (23). Therefore, it remains challenging to determine to what extent existing *Gtf2ird1* exonic mutants model the loss of this gene in WS, where the whole genic locus is deleted. Interestingly, when the whole locus is deleted in the “CD” complete deletion mice, the elevation of *Gtf2ird1* mRNA seen in exonic mutants does not occur (23). However, protein levels appear to stay above 50%, albeit with substantial mouse to mouse variation. Post-mortem patient brain samples, if available, may help to resolve the consequences of the human mutation on GTF2IRD1 protein levels. Further, it may be worth revisiting the consequences of *Gtf2ird1* mutation following generation of full genic deletions in mice. In the meantime, it may be that some of the homozygous point mutations, though different from the heterozygous mutations of WS, may better model WS protein levels as they can result in a reduction of GTF2IRD1 protein (Figure 4). Indeed, most studies of *Gtf2ird1* mutant behavioral consequences have shown atypical phenotypes in homozygous mutant mice (8,9,21).

It is worth noting that even in those mutants with a 50% reduction in protein, transcriptional changes are very subtle, at least at steady state. This is similar to findings in other chromatin modifying knockouts, where final changes in transcription are highly subtle (41–45), even when behavioral consequences can be severe or even lethal, in the case of Mecp2 mutants Guy et al. Nat Gen. 2001 (46). Thus, a second remaining puzzle about these genes is what their role is in regulating transcription at the majority of their binding sites. One possibility is these genes are essential for regulating the proper dynamics of gene expression, something not captured when assessing a population at steady state. Another possibility is they affect phenotypes via actions in rare cell types not easily detected in whole brain RNA-seq. Both hypotheses await further experimentation. Nonetheless, the strong enrichment of ASD and constrained genes among GTF2I and especially GTF2IRD1 targets suggest that these factors may be key regulators. Such functional interactions suggest common pathways across these chromatin-related forms of ASD and intellectual disability.

Regardless of the resiliency at the protein level, we show heterozygous and homozygous mutations of *Gtf2ird1* were sufficient to cause adult behavioral abnormalities. This supports the hypothesis that the N-terminal end of the protein has other important functions beyond DNA binding. Similarly, the N-truncation of GTF2I did not affect DNA-binding genome-wide, but still resulted in behavioral deficits (47). The single *Gtf2ird1* homozygous mutant showed balance deficits, which is consistent across many mouse models of WS. We also observed decreased marble burying. This task is thought to be mediated at least in part by hippocampal function, suggesting a possible disruption of the hippocampus caused by this mutation (48). We also observed an increase in contextual fear response, another cognitive task that is thought to be under hippocampal and amygdalar regulation. Increase in contextual fear was also seen in another *Gtf2ird1* mouse model (49).

Given the prior evidence that these two transcription factors are both involved in cognitive and behavioral phenotypes of WS (7, 50), and the evidence that their shared binding targets regulate synaptic genes, we tested if having both *Gtf2i* and *Gtf2ird1* mutated could modify the phenotype seen when just *Gtf2ird1* was mutated. Contrary to our prediction, we did not see a large effect of adding a *Gtf2i* mutation to differences in transcriptome wide expression or behavioral phenotypes. This was also surprising given that we successfully reduced GTF2I protein and it has been described in the literature as regulating transcription (51). Again, whole E13.5 brain analysis could diminish any effects of transcriptional differences in specific cell types. This potential confound could be overcome using single cell sequencing technologies in the future when those technologies mature and become more reliable for detecting within cell type differences of expression.

When *Gtf2i* was knocked down in the presence of two *Gtf2ird1* mutations, we saw phenotypes in the same direction as the homozygous one base pair insertion *Gtf2ird1* mutant, as well as significant increases in the cued fear memory task. Thus, the behaviors tested in this study seem to be mainly driven by *Gtf2ird1* mutant homozygosity, which is consistent across the different mutations. The one exception seemed to be PPI, in which the knockdown of *Gtf2i* in the presence of two *Gtf2ird1* mutations attenuated the effect of the pre-pulse. There was no effect of homozygous *Gtf2ird1* mutation alone on this phenotype, suggesting that *Gtf2i* and not *Gtf2ird1* is playing a larger role. However, the more severe phenotype was seen in the *Gtf2i*^+/-^ /*Gtf2ird1*^-/-^ mutant compared to *Gtf2i*^+/-^/*Gtf2ird1*^+/-^, suggesting a contribution from both genes. This does not exclude the possibility that *Gtf2i* can modify the phenotype of *Gtf2ird1* knockdown in other behavioral domains. For example, it would be interesting to see the effect of adding a *Gtf2i* mutation on top of a *Gtf2ird1* mutation on social behaviors. However, on the FVB/AntJ background used here and in F1 FVB/AntJ x C57BL/6J crosses used previously, we have not seen any social behavior disruptions when these genes are mutated or the entire WS locus is deleted (23).

Likewise, the fear conditioning effects of even heterozygous *Gtf2ird1* mutation are very clear on the FVB/AntJ background used here, but appear to be masked in F1 FVB x C57BL/6J crosses used in our prior study (23). This indicates that while we did not find evidence for epistasis between *Gtf2i* and Gft2ird1, there is epistasis between these genes and other loci in the genome. Thus, mouse mapping studies using large cohorts of F2 hybrids might provide an opportunity to leverage this strain difference to find genes that interact with *Gtf2ird1* to contribute to these phenotypes. Such studies could help define novel interaction partners for this relatively understudied gene.

Overall, our study has provided the first description of the DNA-binding of both GTF2I and GTF2IRD1 in the developing mouse brain and showed they have unique and overlapping targets. These data will be used to inform downstream studies to understand how these transcription factors interact with the genome. We generated two new mouse models that tested the importance of the N-terminal end of GTF2IRD1 and the effect of mutating both *Gtf2i* and *Gtf2ird1* together. We provided evidence that despite neither gene having much effect on transcription, the *Gtf2ird1* mutation affects balance, marble burying, activity levels and fear memory while adding a *Gtf2i* mutation leads to a larger effect on PPI.

## Materials and Methods

### Generating genome edited mice

We generated *Gtf2i* family mutants as described in (23). To generate unique combinations of gene knockouts, we designed gRNAs targeting early constitutive exons of the mouse *Gtf2i* and *Gtf2ird1* genes. The gRNAs were separately cloned into the pX330 Cas9 expression plasmid (a gift from F. Zhang) and transfected in N2a cells to test for cutting efficiency. DNA was harvested from the cells and cutting was detected using the T7 endonuclease assay. The gRNAs were transcribed in vitro using the MEGAShortScript kit (Ambion, Austin, TX) and the Cas9 mRNA was in vitro transcribed using the mMessageMachine kit (Ambion). The two gRNAs and Cas9 mRNA were injected into FVB/NJ mouse embryos and implanted into donor females. The resulting offspring were genotyped for mutations with gene specific primers designed with the Illumina adapter sequences concatenated to their 3’ prime end to allow for deep sequencing of the amplicons surrounding the expected cut sites. In one line, a large 589 bp deletion in *Gtf2ird1* was detected by amplifying 3.5kb that included exon two, exon three and part of intron three, then using a Nextera library prep (Illumina, San Diego, CA) to deep sequence the amplicon. Here we focus on two founder mice obtained from these injections. Founder lines were bred to FVB/AntJ mice to ensure the mutations existed in the germline and, for double mutant founders, on the same chromosome. The mice were further back-crossed until the mutations were on a complete FVB/AntJ background, which differs from the FVB/NJ background at two loci: Tyr^c-ch^, which gives FVB/AntJ a chinchilla coat color, and the 129P2/OlaHSd wildtype (WT) *Pde6b* allele, which prevents FVB/AntJ from becoming blind in adulthood. Coat color was identified by eye, and the *Pde6b* gene was genotyped by PCR. These mouse lines are available through the MMRRC (accession numbers pending).

### Western blotting

Embryos were harvested on embryonic day 13.5 (E13.5) and the whole brain was dissected in cold PBS then flash frozen in liquid nitrogen. The brains were stored at −80°C until they were lysed. The frozen brain was homogenized in 500ul of 1xRIPA buffer (10mM Tris HCl pH 7.5, 140mM NaCl, 1mM EDTA, 1% Triton X-100, 0.1% DOC, 0.1% SDS, 10mM Na_3_V0_4_, 10mM NaF, 1x protease inhibitor (Roche, Basil, Switzerland)) along with 1:1000 dilution of RNase inhibitors (RNasin (Promega, Madison, WI) and SUPERase In (Thermo Fisher Scientific, Waltham, MA). The homogenate was incubated on ice for 20 minutes then spun at 10,000g for 10 minutes at 4°C to clear the lysate. The lysate was stored in two aliquots of 100μl at −80°C for later protein analysis and 250μl of the lysate was added to 750μl of TRIzol LS and stored at −80°C for later RNA extraction and qPCR. Total protein was quantified using the BCA assay and 25-50μg of protein in 1x Lamelli Buffer with β-mercaptoethanol was loaded onto 4-15% TGX protean gels (Bio-Rad, Hercules, CA). The protein was transferred to a .2μm PVDF membrane by wet transfer. The membrane was blocked with 5% milk in TBST for one hour at room temperature. The membrane was cut at the 75KDa protein marker; the bottom was probed with a GAPDH antibody as an endogenous loading control, while the top was probed with an antibody for either GTF2I or GTF2IRD1. The primary incubation was performed overnight at 4°C. The membrane was washed three times in TBST for five minutes, then incubated with an HRP conjugated secondary antibody diluted in 5% milk in TBST for one hour at room temperature. The blot was washed three times with TBST for five minutes then incubated with Clarity Western ECL substrate (Bio-Rad) for five minutes. The blot was imaged in a MyECL Imager (Thermo Fisher Scientific). Relative protein abundance was quantified using Fiji (NIH) and normalized to GAPDH levels in a reference WT sample. The antibodies used were: Rabbit anti-GTF2IRD1 (1:500, Novus, NBP1-91973), Mouse anti-GTF2I (1:1000 BD Transduction Laboratories, Lexington, KY, BAP-135), and Mouse anti-GAPDH (1:10,000, Sigma Aldrich, St. Louis, MO G8795), HRP-conjugated Goat anti Rabbit-IgG (1:2000, Sigma Aldrich, AP307P) and HRP-conjugated Goat anti-Mouse IgG (1:2000, Bio-Rad, 1706516).

### Transcript abundance using RT-qPCR

RNA was extracted from TRIzol LS using the Zymo Clean and Concentrator-5 kit with on column DNase-I digestion following the manufacturer’s instructions. The RNA was eluted in 30μl of RNase free water and quantified using a Nanodrop 2000 (Thermo Fisher Scientific). One microgram of RNA was transcribed into cDNA using the qScript cDNA synthesis kit (Quanta Biosciences, Beverly, MA). Half a microliter of cDNA was used in a 10μl PCR reaction with 500nM of target specific primers and the PowerUP SYBR green master mix (Applied Biosystems, Foster City, CA). The primers were designed to amplify exons that were constitutively expressed in both *Gtf2i* (exons 25 and 27) and *Gtf2ird1* (exons 8 and 9) and span an intron (23). The RT-qPCR was carried out in a QuantStudio6Flex machine (Applied Biosystems) using the following cycling conditions: 1) 95°C for 20 seconds, 2) 95°C for 1 second, 3) 60°C for 20 seconds, then repeat steps 2 and 3 40 times. Each target and sample was run in triplicate technical replicates, with three biological replicates for each genotype. The relative transcript abundance was determined using the delta CT method normalizing to Gapdh.

### ChIP

Chromatin was prepared as described previously (23). Frozen brains were homogenized in 10mL of cross-linking buffer (10mM HEPES pH7.5, 100mM NaCl, 1mM EDTA, 1mM EGTA, 1% Formaldehyde (Sigma Aldrich)). The homogenate was spun down and resuspended in 5mL of 1x L1 buffer (50mM HEPES pH 7.5, 140 mM NaCl, 1mM EDTA, 1mM EGTA, 0.25% Triton X-100, 0.5% NP40, 10.0% glycerol, 1mM BGP (Sigma Aldrich), 1x Na Butyrate (Millipore, Burlington, MA), 20mM NaF, 1x protease inhibitor (Roche)) to release the nuclei. The nuclei were spun down and resuspended in 5mL of L2 buffer (10mM Tris-HCl pH 8.0, 200mM NaCl, 1mM BGP, 1x Na Butyrate, 20mM NaF, 1x protease inhibitor) and rocked at room temperature for five minutes. The nuclei were spun down and resuspended in 950μl of buffer L3 (10mM Tris-HCl pH 8.0, 1mM EDTA, 1mM EGTA, 0.3% SDS, 1mM BGP, 1x Na Butyrate, 20mM NaF, 1x protease inhibitor) and sonicated to a fragment size of 100-500bp in a Covaris E220 focused-ultrasonicator with 5% duty factor, 140 PIP, and 200cbp. The sonicated chromatin was diluted with 950ul of L3 buffer and 950μl of 3x Covaris buffer (20mM Tris-HCl pH 8.0, 3.0% Triton X-100, 450mM NaCl, 3mM EDTA). The diluted chromatin was pre-cleared using 15μl of protein G coated streptavidin magnetic beads (Thermo Fisher Scientific) for two hours at 4°C. For IP, 15μl of protein G coated streptavidin beads were conjugated to either 10μl of GTF2IRD1 antibody (Rb anti-GTF2IRD1, NBP1-91973 LOT:R40410) or 10μl of GTF2I antibody (Rb anti-GTF2I, Bethyl Laboratories, Montgomery, TX, A301-330A) for one hour at room temperature. 80μl of the pre-cleared lysate was saved for an input sample. 400μl of the pre-cleared lysate was added to the beads and incubated overnight at 4°C. The IP sample was then washed twice with low salt wash buffer (10mM Tris-HCl pH 8.0, 2mM EDTA, 150mM NaCl, 1.0% Triton X-100, 0.1% SDS), twice with a high salt buffer (10mM Trish-HCl pH 8.0, 2mM EDTA, 500mM NaCl, 1.0% Triton X-100, 0.1% SDS), twice with LiCl wash buffer (10mM Tris-HCl pH 8.0, 1mM EDTA, 250mM LiCl (Sigma Aldrich), 0.5% NaDeoxycholate, 1.0% NP40), and once with TE buffer (10mM Tris-HCl pH 8.0, 1mM EDTA). The DNA was eluted from the beads with 200μl of 1x TE and 1% SDS by incubating at 65°C in an Eppendorf R thermomixer shaking at 1400rpm. The DNA was de-crosslinked by incubating at 65°C for 15 hours in a thermocycler. RNA was removed by incubating with 10ug of RNase A (Invitrogen, Carlsbad, CA) at 37°C for 30 minutes and then treated with 140ug of Proteinase K (NEB, Ipswhich, MA) incubating at 55°C in a thermomixer mixing at 900rpm for two hours. The DNA was extracted with 200μl of phenol/chloroform/isoamyl alcohol (Ambion) and cleaned up using the Qiagen PCR purification kit then eluted in 60ul of elution buffer. Concentration was assessed using the high sensitivity DNA kit for Qubit quantification (Thermo Fisher Scientific).

### ChIP-qPCR

Primers were designed to amplify the upstream regulatory element of *Gtf2ird1*. Two off target primers were designed: one 10kb upstream of the transcription start site of *Bdnf* and the other 7kb upstream of the *Pcbp3* transcription start site. The input sample was diluted 1:3, 1:30, and 1:300 to create a standard curve for each primer set and sample. Each standard, input, and IP sample for each primer set was performed in triplicate in 10μl reactions using the PowerUP SYBR green master mix (Applied Biosystems) and 250nM of forward and reverse primers. The reactions were performed in a QuantStudio6Flex machine (Applied Biosystems) with the following cycling conditions: 1) 50°C for 2 minutes, 2) 95°C for 10 minutes, 3) 95°C 15 seconds, 4) 60°C for 1 minute, then repeat steps 3-4 40 times. The relative concentration of the input and IP samples were determined from the standard curve for each primer set. Enrichment of the IP samples was determined by dividing the on target upstream regulatory element relative concentration by the off target relative concentration.

### ChIP-seq

ChIP seq libraries were prepared using the Swift Accel-NGS 2S plus DNA library prep kits with dual indexing (Swift Biosciences, Ann Arbor, MI). The final libraries were enriched by 13 cycles of PCR. The libraries were sequenced by the Genome Technology Access Center at Washington University School of Medicine on a HiSeq3000 producing 1×50 reads.

Raw reads were trimmed of adapter sequences and bases with a quality score below 25 using the Trimmomatic Software (52). The trimmed reads were aligned to the mm10 genome using the default settings of bowtie2 (53). Reads with a mapping quality of less than 10 were removed. Picard tools was used to remove duplicates from the filtered reads (http://broadinstitute.github.io/picard). Macs2 was used to call peaks on the WT IP, *Gtf2ird1*^-/-^ IP, and *Gtf2i*^-/-^/*Gtf2ird1*^-/-^ IP samples with the corresponding sample’s input as the control for each biological replicate (54). Macs2 used an FDR of 0.01 as the threshold to call a significant peak. High confidence peaks were those peaks that had some overlap within each biological replicate for each genotype using bedtools intersect (55). The read coverage for the WT high confidence peaks was determined using bedtools coverage for all genotypes. To identify peaks with differential coverage, we used EdgeR to compare the WT peak coverage files to the corresponding mutant peak coverage; differential peaks were defined as having an FDR < 0.1 (56). To determine GTF2I high confidence peaks, we used peaks that had overlap between all four biological replicates and WT peaks with an FDR < 0.1 and log2FC > 0 when compared to the *Gtf2i*^-/-^/*Gtf2ird1*^-/-^ IP coverage, since this mutation represents a full knockout of the protein.

Annotations of peaks and motif analysis was performed using the HOMER software on the high confidence peaks (57). Peaks were annotated at the transcription start (TSS) of genes if the peak overlapped the +2.5kbp or −1kbp of the TSS using a custom R script. GO analysis on the ChIP target genes was performed using the goseq R package. Comparison to ASD genes entailed testing for overlap between ChIP target genes and the genes identified by (28), their **table S4**, using Fisher’s exact test and assuming 22,007 protein coding genes in the genome based on current NCBI annotation. Slightly more significant results were obtained when replicating this analysis using the SFARI gene database (58), accessed 8/4/2019, and testing for enrichment of score 1 and 2 ASD genes. We also analyzed pLI scores, downloaded from Gnomad (31) on 9/16/2019, of ChIP peaks. Similar results were obtained using pLI cutoffs of .9 and .99. For epigenetic overlaps, we used E13.5 H3K4me3 and E13.5 H3K27me3 forebrain narrow bed peak files from the mouse ENCODE project to overlap with our peak datasets (59). Deeptools was used to generate bigwig files normalized to the library size for each sample by splitting the genome into 50bp overlapping bins (60). Deeptools was used to visualize the ChIP-seq coverage within the H3K4me3 and H3K27me3 peak regions. The LICR TFBS E14.5 whole brain CTCF peak data set was downloaded from the UCSC genome browser and lifted over to mm10 coordinates. TAD analysis of E14.5 cortical neuron HiC data (61) was carried out using domains previously called by arrowhead (Rao et al. Cell 2014, Clemens et al. *Mol Cell* in press). Domain boundaries were defined as 10kb regions centered on the start and end of each domain and lifted over to mm10 coordinates. Overlaps between GTF2IRD1 and GTF2I peaks and other peak datasets was performed using the bedtools fisher function. PhyloP scores for the region underneath the WT ChIP-seq peaks and random genomic regions of the same length were retrieved using the UCSC table browser 60 Vertebrate Conservation PhyloP table. The Epigenome browser was used to visualize the ChIP-seq data as tracks (62).

### RNA-seq

One microgram of E13.5 whole brain total RNA extracted from TRIzol LS was used as input for rRNA depletion using the NEBNext rRNA Depletion Kit (Human/Mouse/Rat). The rRNA-depleted RNA was used as input for library construction using the NEBNext Ultra II RNA library prep kit for Illumina. The final libraries were indexed and enriched by PCR using the following thermocycler conditions: 1) 98°C for 30 seconds, 2) 98°C 10 seconds, 3) 65°C 75 seconds, 4) 65°C 5 minutes, 5) hold at 4°C, repeating steps 2-3 six times. The libraries were sequenced by the Genome Technology Access Center at Washington University School of Medicine on a HiSeq3000 producing 1×50 reads.

### RNA-seq analysis

The raw RNA-seq reads were trimmed of Illumina adapters and bases with quality scores less than 25 using Trimmomatic Software. The trimmed reads were aligned to the mm10 mouse genome using the default parameters of STARv2.6.1b (63). We used HTSeq-count to determine the read counts for features using the Ensembl GRCm38 version 93 gtf file (64). Differential gene expression analysis was done using EdgeR. We compared the expression of genes that are targets of either GTF2IRD1 or GTF2I to non-bound genes by generating a cumulative distribution plot of the average log CPM of the genes between genotypes. GO analysis was performed using the goseq R package.

### Data Availability

All RNA-seq and ChIP-seq data is available at GEO accession GSE138234.

### Behavioral tasks

All animal testing was done with approval from the Washington University in St. Louis Institutional Animal Care and Use Committee. Mice were group housed in same-sex, mixed-genotype cages with two to five mice per cage in standard mouse cages (dimensions 28.5 x 17.5 x 12 cm) on corn cob bedding. Mice had ad libitum access to food and water and followed a 12-hour light-dark cycle (light from 6:00am-6:00pm). Animal housing rooms were kept at 20-22°C with a relative humidity of 50%. All mice were maintained on the FVB/AntJ (65) background from Jackson Labs. All behaviors were done in adulthood between ages P58-P133. A week prior to behavioral testing mice were handled by the experimenter for habituation. On testing days the mice were moved to habituate to the testing room and the experimenter if male for 30 minutes before testing started.

#### Ledge

To test balance, we measured how long a mouse could balance on an acrylic ledge with a width of 0.5cm and a height of 38cm as described (65). The time when the mouse fell off the ledge was recorded up to 60 seconds when the trial was stopped. If the mouse fell off within five seconds, the time was restarted and the mouse received another attempt. If the mouse fell off within the first five seconds on the third attempt, that time was recorded. We tested all mice on the ledge twice, with a rest period of at least 20 minutes between trials. Trial average was used for analysis.

#### One-hour locomotor activity

We assessed activity levels in a one-hour locomotor task, as previously described (65). Mice were placed in the center of a standard rat-sized cage (dimensions 47.6 x 25.4 x 20.6cm). The rat-sized cage was located inside a sound-attenuating box with white light at 24 lux. The mice could freely explore the cage for one hour. An acrylic lid with air holes was placed on top of the cage to prevent mice from jumping out. The position and horizontal movement of the mice was tracked using ANY-maze software (Stoelting Co., Wood dale, IL: RRID: SCR_014289). The apparatus was divided into two zones: a 33 x 11cm center zone and an edge zone of 5.5cm that bordered the cage. The animal was considered in a zone if 80% of the mouse was detected in that zone. ANY-maze recorded the time, distance, and number of entries into each zone. After the task, the mouse was returned to its home cage and the apparatus was thoroughly cleaned with 70% ethanol.

#### Marble burying

Marble burying is a species-specific task that measures the compulsive digging behavior of mice. Normal hippocampal function is thought to be required for normal phenotypes in this task. We tested marble burying as described previously (65). A rat cage was filled with aspen bedding to a depth of 3cm and placed in a sound-attenuating box with white light set to 24 lux. Twenty marbles were placed on top of the bedding in a 5 x 4 evenly spaced grid. The experimental mouse was placed in the center of the chamber and allowed to freely explore and dig for 30 minutes. An acrylic lid with air holes was placed on top of the cage to prevent mice from escaping. After 30 minutes, the animal was returned to its home cage. Two scorers counted the number of marbles not buried (less than two-thirds of the marble was covered with bedding). The number of marbles buried was then determined, and the average of the two scores was used in the analysis. After the marbles were counted, the bedding was disposed of and the cage and marbles were cleaned with 70% ethanol.

#### Pre-pulse inhibition

Pre-pulse inhibition (PPI) measures the suppression of the acoustic startle reflex when an animal is presented with a smaller stimulus (pre-pulse) before a more intense stimulus (startle). Acoustic startle response and pre-pulse inhibition were measured as previously described (23). Startle Monitor II software was used to run the protocol in a sound-attenuating chamber (Kinder Scientific, LLC, Poway, CA). Mice were secured in a small enclosure and placed on a force-sensitive plate inside the chamber. Animals were acclimated to the test chamber for 5 minutes before trials began. Startle amplitude was recorded with 1ms force readings during 65 pseudo-randomized trials that alternated between various stimulus conditions. An auditory stimulus of 120dB was presented for 40ms to illicit the startle response. Pre-pulse inhibition was measured in three different types of trials where the 120dB stimulus was preceded by pre-pulses of 4, 8, and 16 dB above the background sound level (65dB). PPI was calculated by using the average startle response (measured in Newtons) of the 120dB stimulus trials minus the average startle response to the stimulus after each respective pre-pulse level. This was divided by the 120dB stimulus startle response and multiplied by 100 to get a percent inhibition of startle.

#### Contextual and Cued Fear Conditioning

Learning and memory were tested using the contextual and cued fear conditioning paradigm as previously described (66). Contextual fear memory is thought to be driven by hippocampal functioning whereas cued fear is thought to be driven by amygdala functioning. On day one of the experiment, animals were placed in an acrylic chamber (26cm x 18cm x 18cm; Med Associates Inc., Fairfax, VT) with a metal grid floor and a peppermint odor from an unobtainable source. The chamber light was on for the duration of the five-minute task. During the first two minutes, the animal freely explored the apparatus to measure baseline freezing. At 100 seconds, 160 seconds, and 220 seconds, an 80dB white noise tone (conditioned stimulus, CS) was played for a duration of 20 seconds. During the last two seconds of this tone, the mice received a 1.0mA foot shock (unconditioned stimulus, UCS). The animal’s freezing behavior was monitored by FreezeFrame software (Actimetrics, Evanston, IL) in 0.75s intervals. Freezing was defined as no movement besides respiration and was used as a measure of the fear response in mice. After the five-minute task, the mice were returned to their home cage. On day two, we tested contextual fear memory. The mice were placed in the same chamber as day one with the peppermint odor, and freezing behavior was measured over eight minutes. The first two minutes of day two were compared to the first two minutes of day one to test for the acquisition of fear memory. The mice were then returned to their home cage. On day three, to test cued fear, the mice were placed in a new black and white chamber that was partitioned into a triangle shape and had a new coconut scent. The mice were allowed to explore the chamber and the first two minutes were considered baseline. After minute two the 80dB tone (CS) was played for the remaining eight minutes. Freezing behavior was monitored during the entire ten-minute task.

#### Shock sensitivity

We tested the shock sensitivity of the mice to ensure that differences in conditioned fear were not due to differing responses to the shock itself, following a previously established protocol (23). Mice were placed in the apparatus used for day one and day two of the conditioned fear task. The mice were delivered a two second 0.05mA shock and their behavior was observed. The shock was increased by 0.05mA up to 1mA or until the mice exhibited flinching, escape, and vocalization behaviors. Once the mouse had shown each behavior, the test was ended and the final mA used was recorded.

### Statistical Analysis

All statistical analyses were performed in R v3.4.2 and are reported in **Supplemental Table 3**. The ANOVA assumption of normality was assessed using the Shapiro-Wilkes test and manual inspection of qqPlots, and the assumption of equal variances was assessed with Levene’s Test. When appropriate, ANOVA was used to test for main effects and interaction terms. Post hoc analyses were done to compare between genotypes. If the data violated the assumptions of ANOVA, non-parametric tests were performed. If the experiment was performed over time, linear mixed models were used to account for the repeated measures of an animal using the lme4 R package. Post hoc analyses were then conducted to compare between genotypes within time bins. Post hoc analyses were done using the multcomp R package (67). Animals were removed from analysis if they had a value that was 3.29 standard deviations above the mean or had poor video tracking and could not be analyzed.

## Acknowledgements

This work was supported by 1R01MH107515 JDD, and the Autism Science Foundation, and the National Science Foundation Graduate Research Fellowship DGE-1745038 to NDK and DGE-1745038 to KRN. We would like to thank the Genome Technology Access Center for technical support, as well as Dr. Beth Kozel for critical advice on this project. We would also like to thank Dr. David Wozniak and the Animal Behavior Core at the Washington University School of Medicine for their support.

