## Supplemental Figure 1 for "Functions of *Gtf2i* and *Gtf2ird1* in the developing brain: transcription, DNA-binding, and long term behavioral consequences"

A

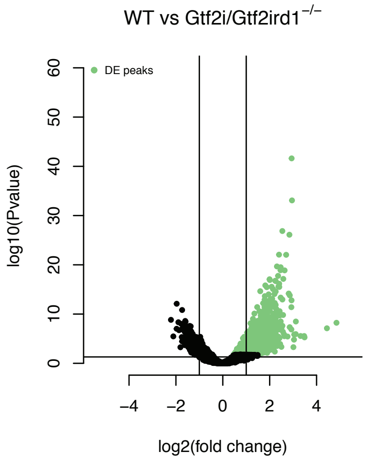


**Supplemental Figure 1: Differential peak binding comparing the WT and homozygous GTF2I IP. A** The highlighted peaks have an FDR < 0.1 and a log2FC > 0. These were used as the high confidence GTF2I peaks.
