## Supplementary figures and images for "Functions of *Gtf2i* and *Gtf2ird1* in the developing brain: transcription, DNA-binding, and long term behavioral consequences"

### Supplemental Figure 2

A

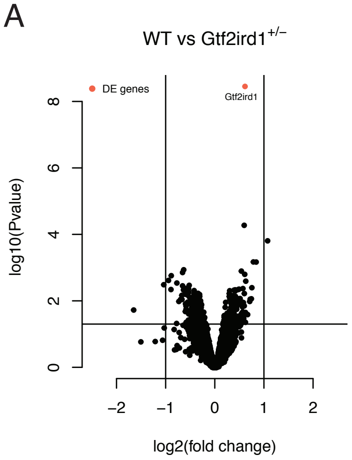


**Supplemental Figure 2: RNA-seq analysis comparing the WT and Gtf2ird1$^{+/-}$. A** Only *Gtf2ird1* showed an FDR < 0.1.
