## Supplemental Figure 3 for "Functions of *Gtf2i* and *Gtf2ird1* in the developing brain: transcription, DNA-binding, and long term behavioral consequences"

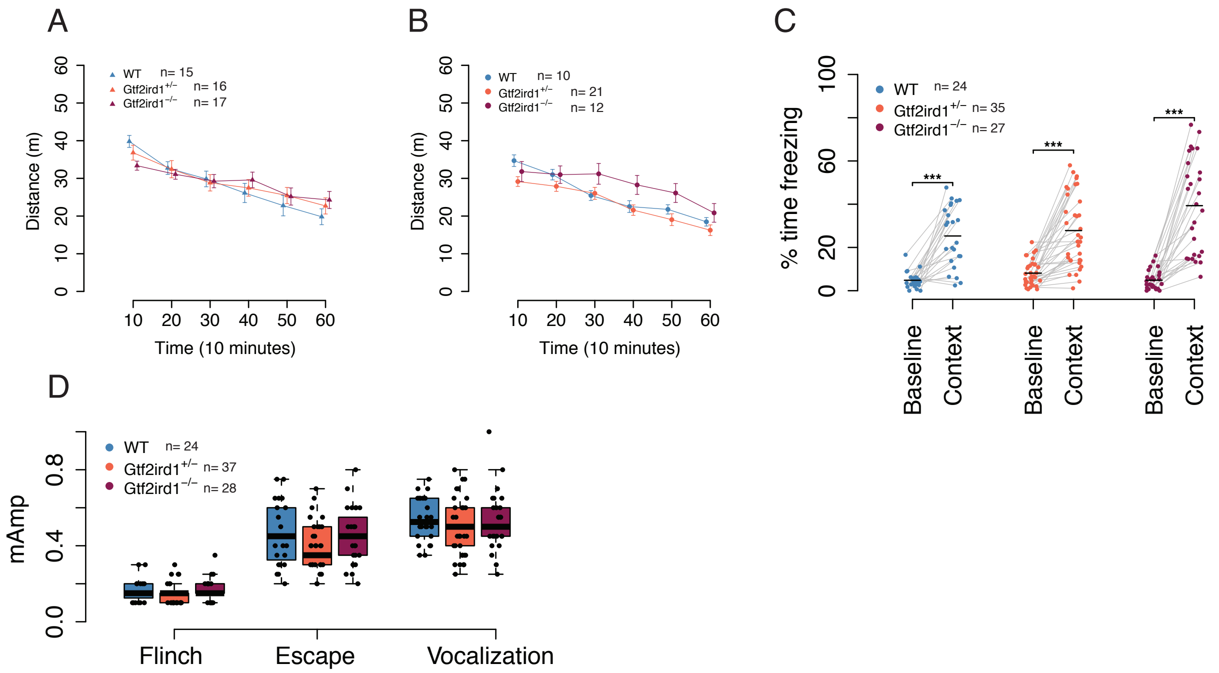


**Supplemental Figure 3: The effects of frameshift mutation in *Gtf2ird1*. A** Female activity levels are similar across genotypes. **B** There is no difference in activity levels among male mice. **C** All genotypes showed a contextual fear response.Baseline refers to the first two minutes of the task on day one and context refers to the first two mintues of the task on day two. **D** There was no difference in shock sensitivity between genotypes.
