## Supplemental Figure 4 for "Functions of *Gtf2i* and *Gtf2ird1* in the developing brain: transcription, DNA-binding, and long term behavioral consequences"

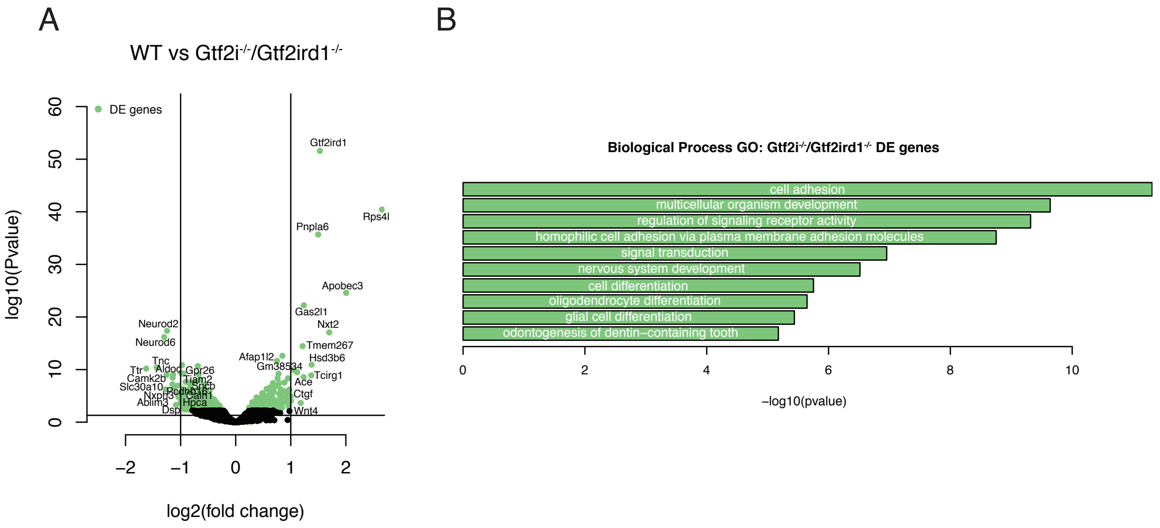


**Supplemental Figure 4: RNA-seq analysis of homozygous double mutant. A** The homozygous double mutant showed significant changes (FDR < 0.1, highlighted in green) across many genes. Genes are labeled that had an FDR < 0.1 and a log2FC > 1 or log2FC < -1. **B** GO analysis of all nominally significant genes.
