## Supplemental Figure 5 for "Functions of *Gtf2i* and *Gtf2ird1* in the developing brain: transcription, DNA-binding, and long term behavioral consequences"

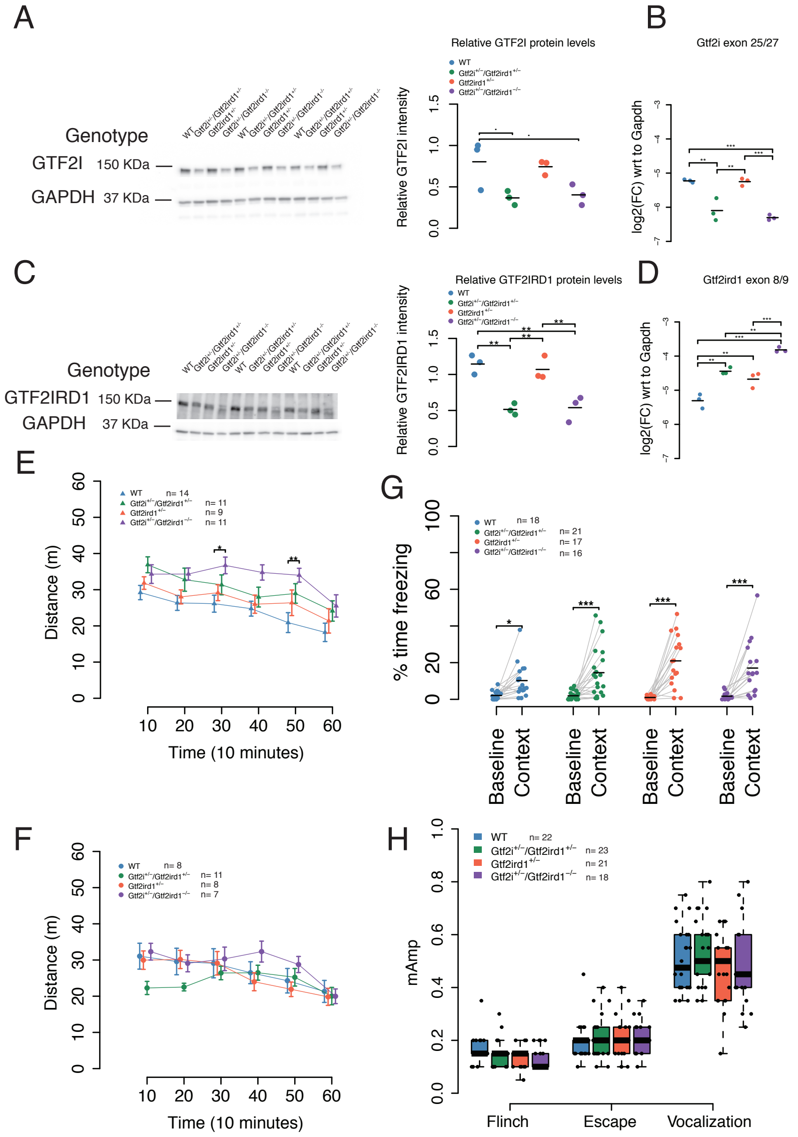


**Supplemental Figure 5: Biochemcial and Behavioral characterization of the Gtf2ird1$^{+/-}$ x Gtf2i$^{+/-}$/Gtf2ird1$^{+/-}$ . A, B** Western blot and qPCR confirm decrease in GTF2I protein and mRNA**. C, D** Western blot shows that the large *Gtf2ird1* deletion decreases the protein, but adding the one base pair insertion mutation does not further decrease the protein made. **D** *Gtf2ird1* mutation increases mRNA abundance. **E** Gtf2i$^{+/-}$/Gtf2ird1$^{-/-}$ females have increased activity levels. **F** Gtf2i$^{+/-}$/Gtf2ird1$^{-/-}$ males to a lesser extent have increased activity. **G** All genotypes showed a contextual fear memory response. **H** There is no difference between genotypes in shock sensitivity.
